# Structural Plasticity and Ligand Promiscuity of CYP3A4 Revealed by Cryo-EM

**DOI:** 10.64898/2026.09.01.748687

**Authors:** Anna Karen Orta, Jan-Hannes Schäfer, Galen J. Correy, Jordan O. Norman, Julius Pampel, Kate K. Huddleston, Hugo MacDermott-Opeskin, Edward B. Miller, Gabriella Reggiano, João P. G. L. M. Rodrigues, Gabriel C. Lander, W. Patrick Walters, James S. Fraser

## Abstract

Cytochrome P450 3A4 (CYP3A4) metabolizes roughly half of all marketed drugs, and its inhibition can cause clinically significant drug-drug interactions. The enzyme accommodates chemically diverse ligands, making binding modes and metabolic outcomes difficult to predict. Previous X-ray crystallography efforts have leveraged a truncated construct without the N-terminal segment that tethers CYP3A4 to the membrane. Here we show that the same construct assembles into a symmetric trimer that can be resolved by cryo-EM and determine structures of both unliganded and ligand-bound CYP3A4. Multiple ligands are resolved with density consistent with several mutually exclusive conformations. Protein remodeling to reshape the binding pocket is concentrated in the F/G loop, which is poorly resolved and unmodeled in many X-ray structures. These features likely underlie the poor predictive performance of co-folding methods on this target. The routine use of cryo-EM to resolve CYP3A4 ligand-bound complexes will provide the ground truth data needed to make predictive models of drug metabolism useful in practice.

## Introduction

Cytochrome P450 enzymes (CYPs) play a central role in the biotransformation of xenobiotics, with CYP3A4 responsible for the metabolism of approximately half of marketed pharmaceuticals (Zanger and Schwab, 2013; Zhang et al., 2024). This widespread activity of CYP3A4 makes accurate prediction of its substrate specificity and inhibition critical for anticipating metabolic liabilities and drug-drug interactions during therapeutic development (Feng et al., 2023; Hayes et al., 2014). The ability of CYP3A4 to accommodate chemically diverse ligands requires structural rearrangements of the active site and access channels, complicating prediction of binding modes and metabolic hotspots (Ekroos and Sjögren, 2006). Many of these rearrangements are typical across all CYPs as part of their heme-dependent monooxygenase catalytic cycle. CYPs catalyze substrate oxidation through a multistep catalytic cycle involving substrate-induced water displacement, sequential electron transfer from cytochrome P450 reductase, formation of an iron-oxo intermediate, and oxygen insertion into the bound substrate (Denisov et al., 2005). Product formation requires precise positioning of the substrate relative to the reactive heme center, thereby linking ligand-binding mode directly to metabolic outcome.

In addition to the catalytic promiscuity of CYP3A4, drug-drug interactions can be additionally complicated by the broad chemical diversity of molecules that can act as inhibitors of its catalytic activity (Dresser et al., 2000). Inhibition of CYP3A4 can greatly extend the pharmacokinetic exposure of co-administered drugs, a property that can be used therapeutically when inhibitors are co-administered with substrates. The most famous example of this co-dosing is nirmatrelvir/ritonavir (Paxlovid), in which ritonavir acts as a strong CYP3A4 inhibitor, thereby hindering the metabolism of nirmatrelvir for the treatment of SARS-CoV-2 (Bege and Borbás, 2024).

CYP3A4 can be inhibited through two mechanistically distinct binding modes (Types I and II) (Ahlström and Zamora, 2008). Type I inhibitors displace a water molecule distal to the heme iron and are dominated by interactions with the protein residues lining the heme-binding pocket. In contrast, Type II inhibitors directly coordinate the heme iron through a nitrogen atom, yielding tight binding and strong inhibition and causing a red-shifted Soret peak (Schenkman et al., 1967). Predicting CYP3A4-ligand binding modes is additionally complicated because structurally related molecules can occupy the active site in distinct orientations, with multiple molecules in the active site, or switch between substrate and inhibitor behavior depending on subtle changes in chemistry or in concentration (Davydov and Halpert, 2008; Ekroos and Sjögren, 2006). Notably, computational approaches have successfully predicted multiple sites of metabolism for certain CYP3A4 substrates, demonstrating that regioselectivity can be captured in specific cases (Yuki et al., 2012). Time-dependent inhibition adds further complexity, as certain compounds may become reactive metabolites that can undergo a second round of oxidation, or inhibit CYP3A4 altogether (Eng et al., 2020). Collectively, multiple binding modes and potential catalytically generated products present a persistent challenge for computational prediction and rational control of CYP3A4-mediated metabolism during drug development.

Structural studies have provided important insights into CYP3A4 function and ligand metabolism. Previous X-ray crystal structures have offered high-resolution views of ligand coordination by leveraging N-terminally truncated constructs that improve solubility and crystallizability by eliminating the anchoring N-terminal transmembrane helix (Sevrioukova and Poulos, 2012, 2010; Yano et al., 2004). However, these structures are enriched for ligands that stabilize crystallizable conformations and ritonavir-derived inhibitors, leaving portions of the chemical space that interact with different conformations of CYP3A4 under-sampled. Several conformational transitions in CYP3A4 regulate substrate access pathways and enable multiple binding orientations within the catalytic cavity (Hayes et al., 2014). In particular, previous studies have noted that helices F and G, the F/G loop, and active-site residues that coordinate the substrate relative to the heme can rearrange depending on the ligand (Chuo et al., 2019). Simulations and solution measurements also suggest that additional, more diverse conformations are populated for some ligand-binding mechanisms (Chuo et al., 2019; Paço et al., 2023). Therefore, incomplete structural coverage may limit efforts to model binding modes for chemically diverse substrates using physics-based and machine learning approaches that rely on a restricted set of receptor conformations (Zhai et al., 2023). A more comprehensive structural view of these important conformational states, including alternate ligand poses and the residues that stabilize them, would strengthen both mechanistic interpretation and computational modeling of CYP3A4-mediated metabolism and its inhibition.

Here, we expand the structural characterization of CYP3A4 using cryo-EM, which can help overcome several limitations of classical X-ray crystallography. We demonstrate that the same soluble construct widely used in biochemical and crystallographic studies forms a trimeric assembly that can be resolved at high resolution. We compare structures of an inhibitor-free state, complexed with inhibitors previously resolved by crystallography (ritonavir, azamulin, and ketoconazole), and a previously unresolved complex with a substrate vardenafil. These cryo-EM reconstructions reveal alternative substrate conformations within the active site, ligand poses distinct from those predicted by co-folding methods, and protein conformational adaptations. By integrating our new data with previous X-ray crystal structures, our work extends the structural framework for understanding CYP3A4 substrate accessibility, inhibitor recognition, and conformational rearrangements, providing experimental ground truth for future computational modeling of CYP3A4-mediated metabolism and enabling more efficient pharmacologic modulation of CYP3A4.

## Results

### The structure of unbound CYP3A4 via cryo-EM reveals a stable trimeric assembly

Most publicly available CYP3A4 structures utilize a soluble cytochrome construct lacking its N-terminal helix, dating back to the construct used for the initial X-ray structure (PDB:1TQN) (Yano et al., 2004). With recent advancements in cryo-EM structure determination, we sought to establish a pipeline to determine the structure of the soluble CYP3A4 construct by cryo-EM, even though its molecular weight is only 55 kDa. The His6-tagged Δ3-22 CYP3A4 construct, herein referred to as CYP3A4, was recombinantly expressed in *E. coli* supplemented with 5-aminolevulinic acid and purified using nickel-affinity resin and size-exclusion chromatography. The size-exclusion chromatogram revealed a predominant higher-molecular-weight peak consistent with a trimeric assembly, along with a shoulder at higher molecular weight suggestive of a tetrameric or even higher-order species (**Supplementary Figure 1**). Fractions were analyzed by SDS-PAGE, revealing pure CYP3A4, and applied to cryo-EM grids for data collection.

Initial datasets exhibited strong preferred orientation, yielding anisotropic maps (**Supplementary Figure 1**). Supplementing the sample with 0.08% DM produced marginal improvement, but overall particle quality was superior in the original dataset. Preferred orientation was mitigated by training a Topaz model on side-view particles, combining these particles with the original top-view particle set, and applying 3D classification (**Supplementary Figure 1**). Final half-maps from Non-Uniform Refinement were subjected to AR-Deconvolution (Li et al., 2025) followed by sharpening with EMReady2 (Cao et al., 2025).

This structure reveals how the Δ3-22 CYP3A4 construct, which is widely used for crystallography and biochemical characterization, can assemble into higher-molecular-weight ensembles that facilitate cryo-EM data processing (**Fig. 1**A). Although monomeric and dimeric states were observed, refinement yielded a trimeric cryo-EM density map at an overall resolution of 3.25 Å. The complex adopts a C3-symmetric arrangement, with protomers engaging through interfaces formed primarily by the N-terminal coil A′ and helix G′ (**Fig. 1**B), resembling a three-leaved flower. Local resolution ranges between 2.9 Å and 4.2 Å, with the best-resolved regions at the CYP3A4 core where the heme cofactor likely aids in particle alignment (**Fig. 1**C). Each protomer closely resembles known CYP3A4 crystal structures, with a backbone RMSD of 0.56 Å to PDB: 1TQN, including the canonical P450 fold and heme-binding pocket (Yano et al., 2004). Displacement is concentrated in several peripheral loops, particularly at the G/H, E/F, D/E, H/I, and I/J regions (**Fig. 1**D), consistent with the inherent conformational flexibility of these regions. Additionally, higher B-factors from earlier X-ray structures (PDB: 1TQN) support localized flexibility within these regions. Helix G of one protomer packs against the A′ loop of its neighbor, forming a predominantly hydrophobic interface that stabilizes the trimer and creates a central hydrophobic core (**Supplementary Figure 2A**). We observe a large density at the core of the trimer, which further non-symmetric refinements demonstrated is not proteinaceous, and is likely a side effect of the high glycerol concentrations required for purification (**Supplementary Figure 2B**).

**Figure 1:**
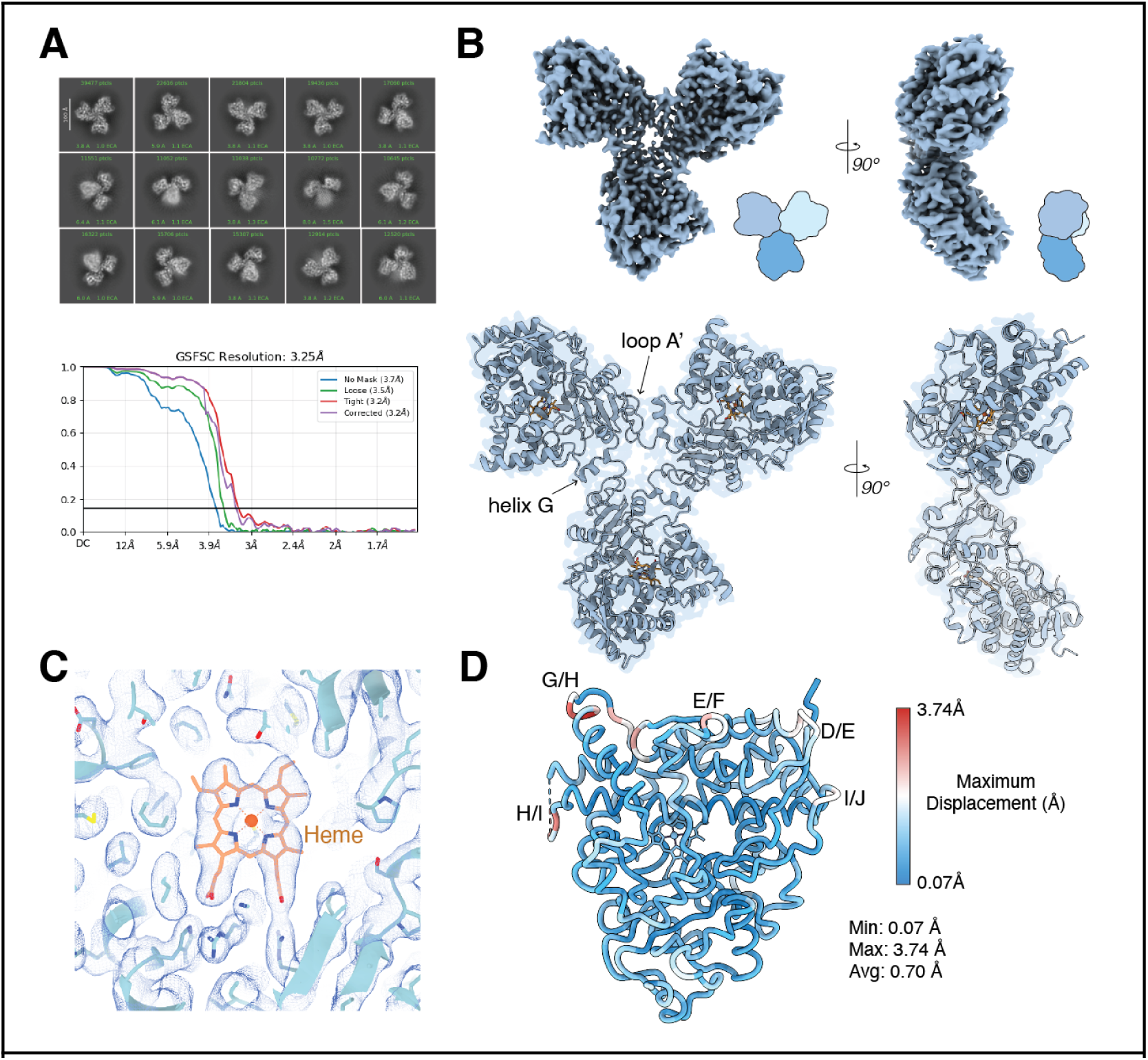
**A.** Representative 2D classes of the final model (top) and corresponding GSFSC curve of the final map to a reported resolution of 3.25 Å at an FSC of 0.143. **B.** Density map of CYP3A4 in a trimeric conformation (top) and model shown in blue (bottom), heme ligand is shown in orange. Insets show a cartoon model of the trimeric assembly colored in shades of blue. **C.** Density at the heme-core and ligand binding site with corresponding side chain densities. **D.** CYP3A4 protomer colored by displacement magnitude relative to PDB:1TQN. The highest displacement in Å is shown in red, lowest displacement in blue. Loops with the highest displacement are labeled.

Although the trimeric assembly observed here lacks the N-terminal transmembrane helix, the hydrophobic oligomerization interface maps to the region where the transmembrane helix would be expected to insert into the membrane (**Supplementary Figure 2C**). In our cryo-EM model, this interface is formed between Leu44 and Pro45 of one protomer and Phe228 and Val235 of the neighboring protomer. The geometry of the trimer is consistent with a model in which the native, membrane-anchored enzyme could adopt a similar oligomeric arrangement. In a membrane context, the transmembrane helices could act to stabilize the interface and the soluble domains could project away from the membrane surface in a concave orientation (**Supplementary Figure 2**). In this hypothetical configuration, the F/G loop remains fully exposed and accessible, suggesting that oligomerization does not occlude substrate entry or active site access. Consistent with this interpretation, the oligomerization interface is spatially remote from the active site and heme-binding cavity, supporting the conclusion that the trimeric state does not compromise the structural integrity or ligand-binding competence of the individual protomers.Ritonavir shows minimal changes relative to X-ray structures

Having established the capabilities of cryo-EM to resolve a novel oligomeric state of CYP3A4, we sought to image previously characterized ligands bound to CYP3A4 to examine any difference relative to the X-ray structures. We started with ritonavir, which is a potent CYP3A4 inhibitor that coordinates the heme iron through its thiazole nitrogen in a canonical Type II binding mode. Ritonavir and its analogs have been extensively characterized by X-ray crystallography (Sevrioukova, 2017; Sevrioukova and Poulos, 2013, 2010). Ritonavir was solubilized and incubated with CYP3A4 at a 10:1 molar ratio with a final concentration of 5% glycerol, 5% DMSO and 2.5% ethanol in phosphate buffer. The sample was incubated for two hours at room temperature to ensure active site saturation and diluted in a 1:1 ratio to phosphate buffer to reduce the glycerol, DMSO and ethanol concentrations immediately prior to vitrification.

Data was collected similarly to the uninhibited structure and while the sample still exhibited some preferred orientation, the drug-bound datasets also contained a larger population of alternate views which mitigated the anisotropy in the final maps. Particles were picked using 2D classes from the unbound dataset as templates, followed by particle curation. Under-represented views from the curated particle stack were then used for Topaz training and picking (**Supplementary Figure 3**), followed by heterogeneous refinement and 3D classification. This map reached a resolution of ∼3.6 Å and was used for model building and refinement. The final map was sharpened with EMReady2, without the need of AR-Deconvolution. The preferred orientation observed in our unbound dataset was likely mitigated by the combination of DMSO and ethanol required for drug solubility.

The high-resolution reconstruction displays unambiguous density for ritonavir within the active site and a well-resolved heme environment (**Fig. 2**). Consistent with prior crystallographic studies, ritonavir adopts a single dominant conformation and coordinates the heme iron directly through its thiazole nitrogen (**Fig. 2**A). Structural comparison with available crystal structures reveals our structure resembles that of PDB:3NXU (Sevrioukova and Poulos, 2010), with an overall Cα RMSD of 0.66 Å, ligand RMSD 1.193 Å, and minimal deviations outside the active site (**Fig. 2**B,C). While the the positions of canonical contact residues lining the I-helix and active site cavity are mostly conserved, and no large-scale helix F/G rearrangements are observed relative to the crystal structure, we observe a distinct state from residues 211-241 comprising of helices F’, G’ and loop F/G (**Fig. 2**B). We also observe some notable ligand-level differences, including a ∼1 Å movement of the valine-like moiety in ritonavir and an associated rotamer shift. The different conformations of the F/G region are consistent with the binding differences between ritonavir analogues to CYP3A4 (**Supplementary Figure 4**). These results confirm that ritonavir enforces a rigid, low-variance active site conformation and highlight the F/G region of flexibility even in tightly-bound ligands like ritonavir. They also reveal that the cryo-EM pipeline can identify subtle conformational differences for ligand-binding features relative to the crystallographic models.

**Figure 2:**
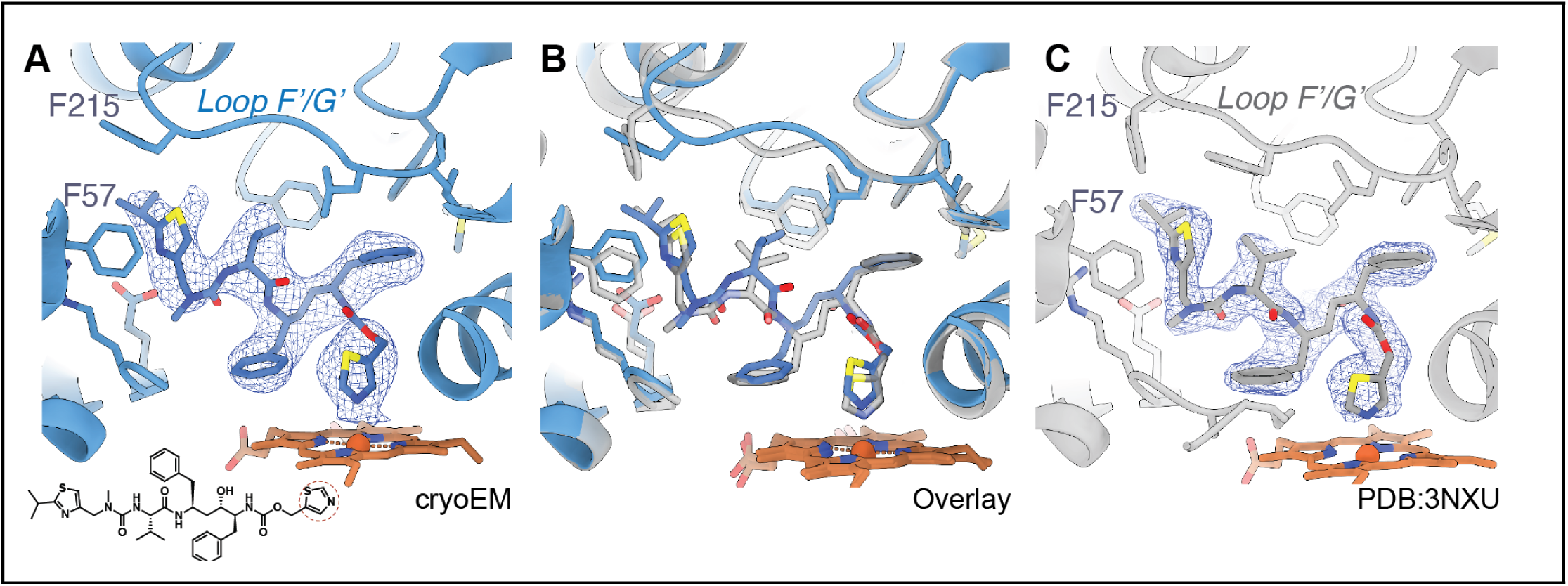
**A.** cryo-EM structure of CYP3A4 bound to ritonavir (blue) along with the corresponding ligand density (mesh). Contact residues are shown as side chains. The chemical structure of ritonavir is shown at the bottom left, with the heme-coordinating moiety highlighted by a red-dotted circle. **B.** Overlay of the crystal structure of ritonavir-bound CYP3A4 (PDB:3NXU) with the lowest RMSD to the model in this work (0.65 Å). **C.** Crystal structure (gray) and corresponding electron density of ritonavir (blue mesh).

### Ketoconazole shows multiple conformations that maintain heme-coordination

Next we wanted to use cryo-EM to resolve a structure where there was ambiguity in binding mode previously observed by crystallography. Ketoconazole is a potent CYP3A4 inhibitor that was previously resolved in two conformations by crystallography (PDB: 2V0M) (Ekroos and Sjögren, 2006) (**Fig. 3**). The primary conformation exhibits a Type II interaction like ritonavir, where the inhibitor engages the heme iron through nitrogen coordination. The secondary conformation is more distant from the heme and is presumably stabilized by packing interactions with the primary conformation (**Fig. 3**C). Following our established cryo-EM preparation procedure we used for ritonavir, we generated a ketoconazole-bound reconstruction to ∼3.8 Å resolution. Although our reconstruction is broadly consistent with the X-ray structure and also reveals heterogeneity in the active site (**Fig. 3**A), the exact details of the cryo-EM density differ. Despite the relatively low global resolution of the EM reconstruction, the density at the active site is sufficiently well resolved to support distinct interpretations, as we explain below.

**Figure 3:**
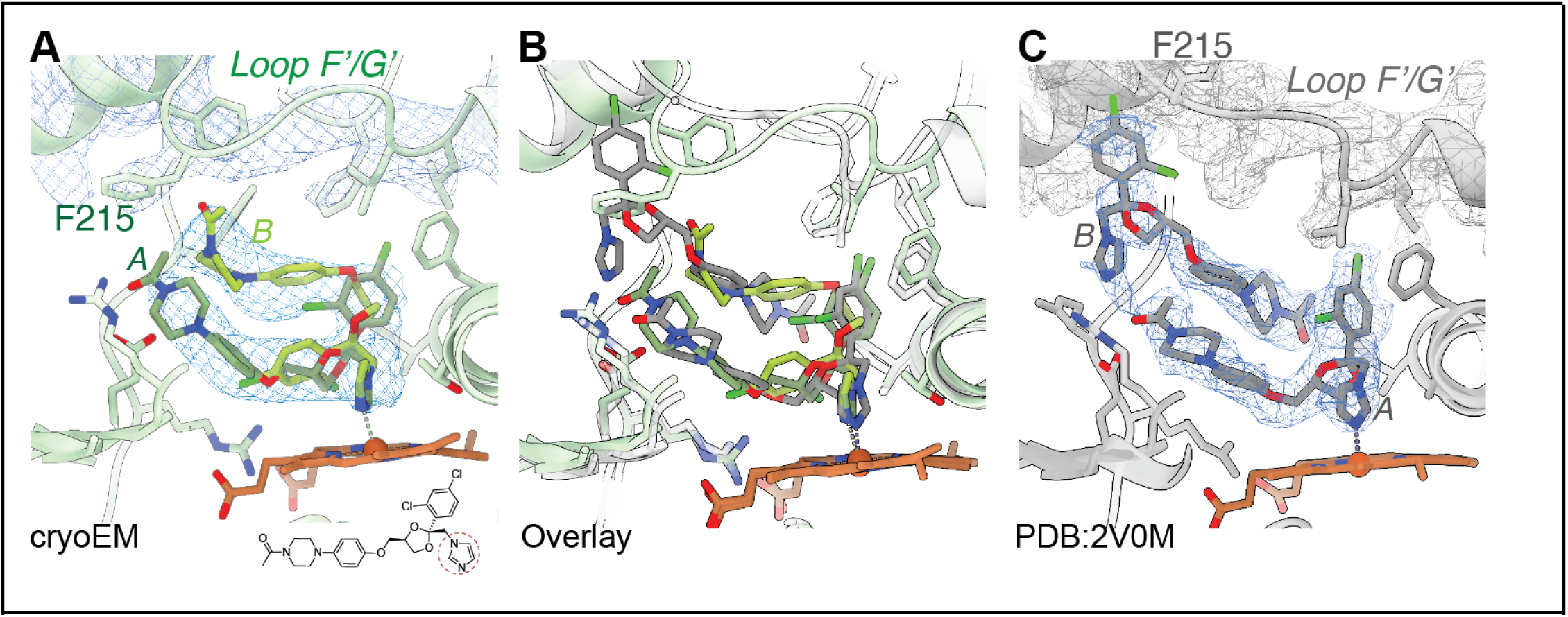
**A**. Cryo-EM structure showing alternate ketoconazole conformations (green) with side chains for contact residues shown, and density as a blue mesh. The chemical structure of ketoconazole is shown in the bottom left, with the heme-coordinating domain highlighted by a red-dotted circle. **B.** Overlay of crystal structure PDB:2V0M (gray) to the cryo-EM structure (green). The reorientation of loop F/G is highlighted. **C**. PDB:2V0M electron density (blue mesh) and model of two stacked ketoconazole poses as seen in the crystal structure.

Relative to the crystal structure, the CYP3A4 protein in the cryo-EM model has an altered orientation of the F/G loop (**Fig. 3**A,B) which adopts a more closed state with the F/G loop constricting the size of the active site compared to the large active site observed in PDB: 2V0M. In the crystal structure, this open conformation accommodates a second ketoconazole molecule stacked against the primary heme-coordinating molecule (**Fig. 3**C). Our cryo-EM reconstruction is, instead, consistent with two mutually exclusive ketoconazole conformations, and cannot physically accommodate two molecules simultaneously (**Fig. 3**A). Both conformations maintain direct imidazole nitrogen coordination to the heme iron and preserve the defining feature of Type II inhibition.

The ketoconazole used in this study is a racemic mixture of the (2S,4R) and (2R,4S) stereoisomers. We hypothesized that these enantiomers bind in distinct states and that are averaged into a single continuous density. To test this, two conformations were refined using Phenix/OPLS with the distance to the coordinating heme iron as a restraint. These models are consistent with the isomers adopting similar conformations of the heme-coordinating imidazole ring, after which the density then branches into the two alternative stereochemical paths. The (2S,4R) ketoconazole molecule was modeled in a favorable conformation within the density closer to the heme, while the (2R,4S) ketoconazole was positioned farther towards the F’/G’ loop. This connected density is consistent with the reported lack of isomer-specificity in ketoconazole binding, as CYP3A4 does not preferentially bind either stereoisomer (Blass et al., 2016). While several hydrophobic residues make contact with both states, the molecule closest to the heme makes unique contacts with Leu210, Phe241, Arg372, Glu374, and Gly481, and the molecule in the distal density uniquely contacts Phe108 and Phe220 in the F/G loop. These results indicate that both ketoconazole stereoisomers engage the heme using the imidazole ring while using distinct hydrophobic contacts to stabilize alternate binding poses.

While this racemate arrangement has some spatial overlap with the X-ray conformations (**Fig. 3**B), the mutually exclusive binding observed by EM is distinct from the simultaneous packing arrangement modeled from crystallography. To confirm the placement of the second molecule in the X-ray model, we deleted it and re-refined the structure using simulated annealing to reduce the bias from the modeled conformation. The resulting omit maps are consistent with the deposited model where two molecules pack against each other and inconsistent with two mutually exclusive alternative conformations (**Supplementary Figure 5**). Collectively, these results demonstrate key differences between the two techniques, with mutually exclusive conformations revealed by cryo-EM and simultaneous binding of stacked molecules revealed by X-ray crystallography.

### Azamulin also occupies multiple conformations

While ritonavir and ketoconazole are Type II inhibitors that directly coordinate the heme iron, many other inhibitors are Type I and do not make this key contact. To resolve a Type I inhibitor by Cryo-EM, we selected azamulin, which has relatively high IC_50_ (0.026-0.24 µM) to CYP3A4 and like ketoconazole has some ambiguity in its previously described binding modes resolved by X-ray crystallography (Ekroos and Sjögren, 2006; Sevrioukova, 2019; Stresser et al., 2004). Characteristic of Type I inhibitors, azamulin does not directly coordinate the heme iron, instead its pleuromutilin ring displaces the heme-coordinating water molecule without forming a new coordinating contact to the heme. Prior crystal structures of CYP3A4 (PDB: 6OOA) and CYP3A5 (PDB: 7SV2) bound to azamulin identified a binding mode in which the pleuromutilin domain is presented toward the heme iron, and molecular dynamics simulations corroborated hydroxyl coordination specifically by residue Ser119 (Hsu and Johnson, 2022; Liu et al., 2025; Sevrioukova, 2019). In addition, the CYP3A5 structure has a secondary “stacked” azamulin molecule, resembling the similar arrangement in CYP3A4 bound to ketoconazole.

In our cryo-EM reconstruction of CYP3A4 bound to azamulin, we observe two key differences in the protein backbone compared to the unbound state. First, the region spanning the E loop through the G helix is displaced with increased disorder in the F/G loop. Second, the C-terminal loop is also displaced. Together, these features combine to create a larger active-site cavity (**Fig. 4**A). Within this cavity, our cryo-EM reconstruction resolves two mutually exclusive azamulin conformations, both of which place the pleuromutilin ring system above the heme (**Supplementary Figure 6**). In both conformations, the hydroxyl group is coordinated by S119 and the pleuromutilin ring is surrounded by several hydrophobic residues, although the ring flipped around the axis defined by the hydroxyl (**Fig. 4**B). The first conformation closely resembles the azamulin-bound CYP3A4 and CYP3A5 crystal structures (**Fig. 4**C,D). The second conformation is inconsistent with a stacked arrangement and interpreted as a mutually exclusive pose. In this secondary conformation, the pleuromutilin ring rotates to present a distinct face of the molecule toward the heme iron, while the triazole moiety is positioned towards the space occupied by the stacked conformation in CYP3A5. Non-protein densities colocalizing with previously identified water-molecule positions are observed above the second triazole conformation, suggesting that ordered solvent is retained in this pose and may influence the geometry of the encounter complex relevant to mechanism-based inactivation. The densities observed in this study also indicate the extent of solvent exposure of the active site in the azamulin-bound state, with various unmodeled and continuous densities from within the active site towards the F/G and C-terminal loop entry site (**Fig. 4**B). This flexibility underlies the fact that the pleuromutilin ring can adopt various conformations above the heme, and that the triazole moiety can engage residues on distinct faces of the active site while anchored by interactions with S119. Collectively, our cryo-EM data show the dynamic nature of CYP3A4 as it adapts to azamulin as a Type I inhibitor.

**Figure 4:**
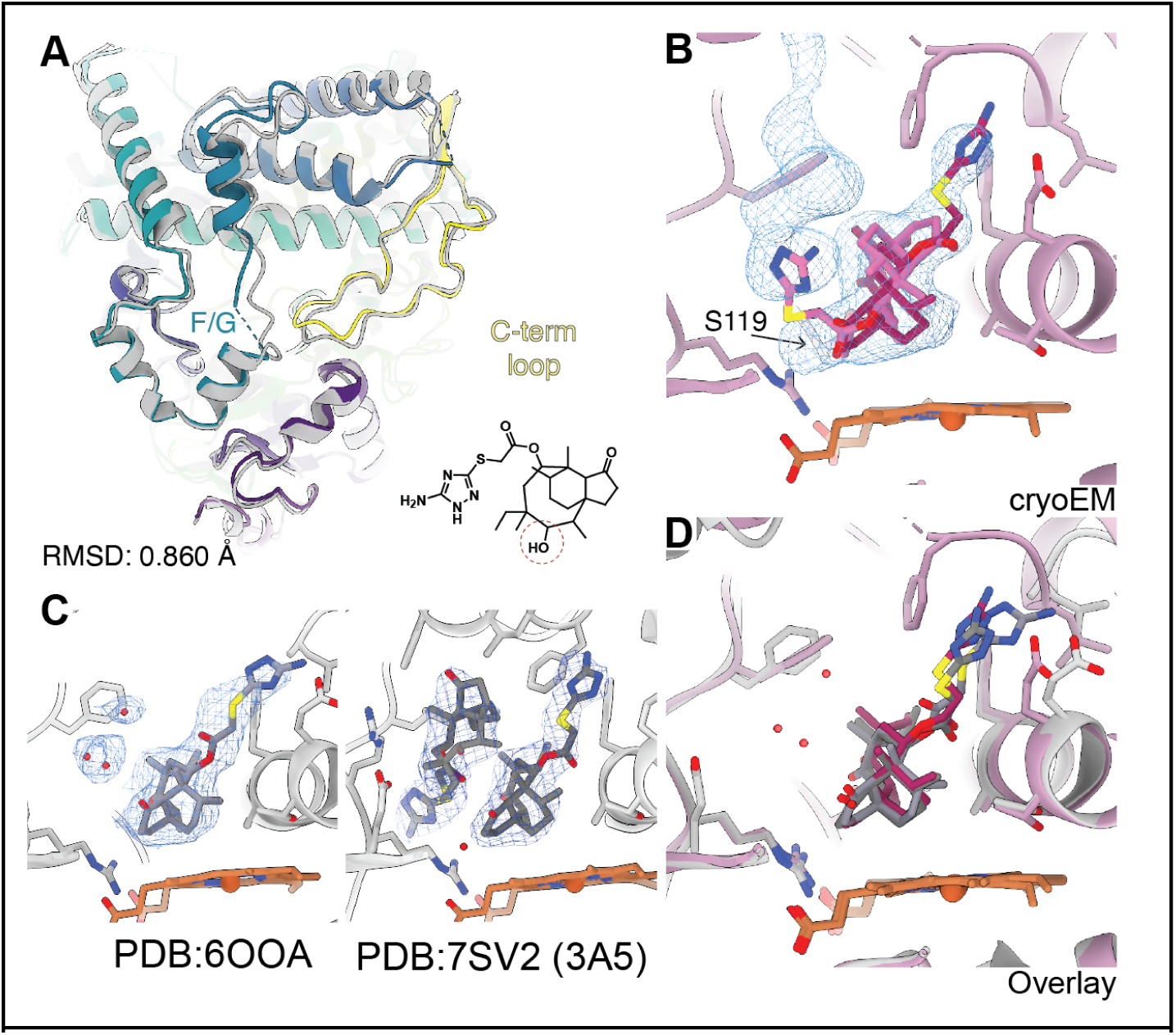
**A.** Viridis-colored structure of CYP3A4-azamulin overlaid against the unbound structure (gray). Loops where differences were observed between the two structures are labeled as F/G for F helix, G helix and the connecting loop, as well as the C-terminal loop. **B.** Azamulin molecule modeled into the density (blue mesh) in two mutually exclusive states (magenta) **C.** PDB: 6OOA (left) and 7SV2 (right) models of azamulin-bound CYP3A4 and CYP3A5, respectively. **D.** Overlay of the common state in all three structures. Contact residues to our model, along with the model of those side chains in the crystal structure of CYP3A4 (6OOA, light gray) are shown. Water molecules from both structures are shown as red dots.

### Structure of CYP3A4 bound to vardenafil reveals multiple conformations

Next, we sought to apply these methods to a ligand without an available structure in the PDB. We selected vardenafil, a PDE5 inhibitor and known CYP3A4 substrate (Ku et al., 2008)whose N-desethylated metabolite acts as a 1.4 μM CYP3A4 inhibitor (DailyMed, 2003). We selected vardenafil because co-folding methods such as Boltz-2 (Passaro et al., 2025) did not converge on a single dominant binding mode (**Supplementary Figure 7**). Some of those binding modes indicated the direct coordination of the heme iron by the non-bridgehead nitrogen of the fused imidazole ring (Type II binding mode), whereas others predict that the piperazine ring sits above the heme without direct coordination (Type I binding mode). To resolve this, we measured absorbance spectra in the presence of each ligand (**Fig. 5**A). Vardenafil produced neither the red-shifted Soret peak characteristic of Type II coordination, observed here with ritonavir, nor the high-spin feature near 390 nm characteristic of Type I binding, observed with azamulin. The near-absence of any spin-state perturbation induced by vardenafil argues against direct coordination of the heme iron.

**Figure 5:**
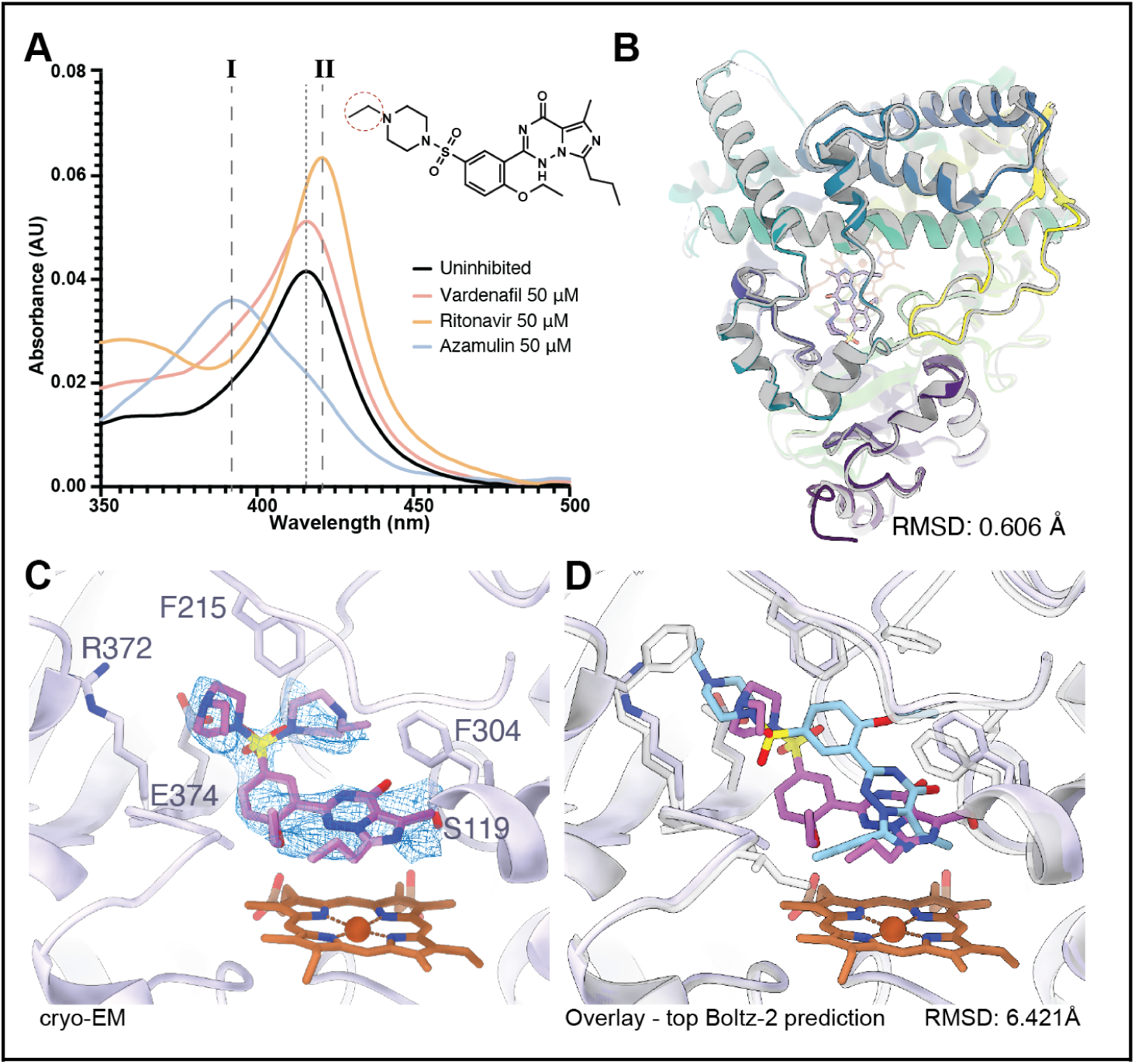
**A**. Soret peak absorbance experiment of CYP3A4 bound to a Type I (azamulin, blue) and Type II (ritonavir, orange) inhibitor compared to vardenafil (pink). The unbound state is shown in black. The structure of vardenafil is shown on the top right corner, with the region that undergoes desethylation highlighted in a dotted circle. **B**. Structural alignment of vardenafil-bound CYP3A4 (viridis) to the unbound state (gray) showing a Cα RMSD: 0.61 Å. **C**. Model of two alternate poses of vardenafil (purple) into cryo-EM density (blue mesh). **D**. Cryo-EM model overlayed with the top scored Boltz-2 prediction showing coordination of the heme group through the same functional group.

The cryo-EM reconstruction of the CYP3A4-vardenafil complex refined to ∼3.2 Å resolution, with minimal changes relative to the unliganded structure (**Fig. 5**B). The resolution was sufficient to unambiguously position the fused imidazole ring above the heme iron (**Fig. 5**C). However, the density was weaker for the piperazine ring and consistent with two alternative poses, neither of which implicated the metabolized ethyl extension in critical interactions (**Fig. 5**C). Interestingly, these conformations do not match the top-ranked predictions from Boltz-2 (**Fig. 5**D, **Supplementary Figure 7**). The Boltz-2 top-scored model places the piperazine ring in a geometry inconsistent with the observed density, with ligand RMSD of 5.43 Å to conformer A and 4.08 Å to conformer B of the cryo-EM model **Fig. 5**D). Collectively, these results demonstrate that experimentally determined cryo-EM structures can resolve binding modes inaccessible to current predictive algorithms.

### Structure prediction methods have a high failure rate on CYP3A4

The discordance between the vardenafil Boltz-2 predictions and the experimentally determined structure (ligand RMSD > 4 Å) led us to question whether such methods generally perform poorly for CYP3A4, given its evolutionary history as a promiscuous xenobiotic metabolizer. To test this idea, we predicted the complexes for all 109 deposited structures in the PDB with Boltz-2 (Passaro et al., 2025). Of these, 89 were determined prior to the training date cutoff for Boltz-2 and are therefore “re-predictions” of complexes the model could have memorized. Even for these 89 predictions, the success rate is 61%, with a successful prediction being defined as having a ligand RMSD ≤2 Å (**Fig. 6**A). This success rate is much lower than the reported PDB-wide rate (Škrinjar et al., 2026) and even lower than the prospective tests on recent Mac1 ligands (Kim et al., 2025). The performance of Boltz-2 on newer PDB complexes not in the training set was significantly lower still, at 40% (**Fig. 6**A). Of these newer complexes, notable successes not in the training set include PDB: 9COT (1.09 Å RMSD) and PDB: 9BVC (1.13 Å RMSD), both of which feature an aromatic nitrogen complexed with the heme iron in a Type II binding mode (**Fig. 6**B). However, the similarities of both complexes to the training set differ substantially with PDB: 9COT having a maximum Tanimoto value of 0.76 (above the test set average of 0.40), while PDB: 9BVC sits at 0.16, which suggests that Type II binding may guide correct placement even when chemical similarity is low. However, overall, there was no obvious relationship between similarity to the training set and accuracy of prediction (**Supplementary Figure 8**). Failures are dominated by missed interactions or ligand placement problems (**Fig. 6**C). The most egregious failures occurred in complexes where progesterone, which is both a substrate and an allosteric binder, was bound in an allosteric pocket (PDB: 1W0F, 5A1R and 5A1P). In these cases, co-folding placed the ligand near the heme, more than 15 Å from the allosteric binding pocket. This aligns with recent results that conclude that co-folding methods struggle to position allosteric ligands (Nittinger et al., 2025). Collectively, these results suggest that co-folding methods may struggle with promiscuous targets like CYP3A4 and other avoid-ome targets (e.g. PXR, P-gp, etc) in particular (Fraser et al., 2026). The origins of this difficulty may lie in the promiscuity of the binding site, the heme cofactor, or the vast chemical space accessible to this protein. As methods for fine-tuning co-folding models become widely available, increased structural coverage of ligands may improve the co-folding performance for CYP3A4 and related targets.

**Figure 6:**
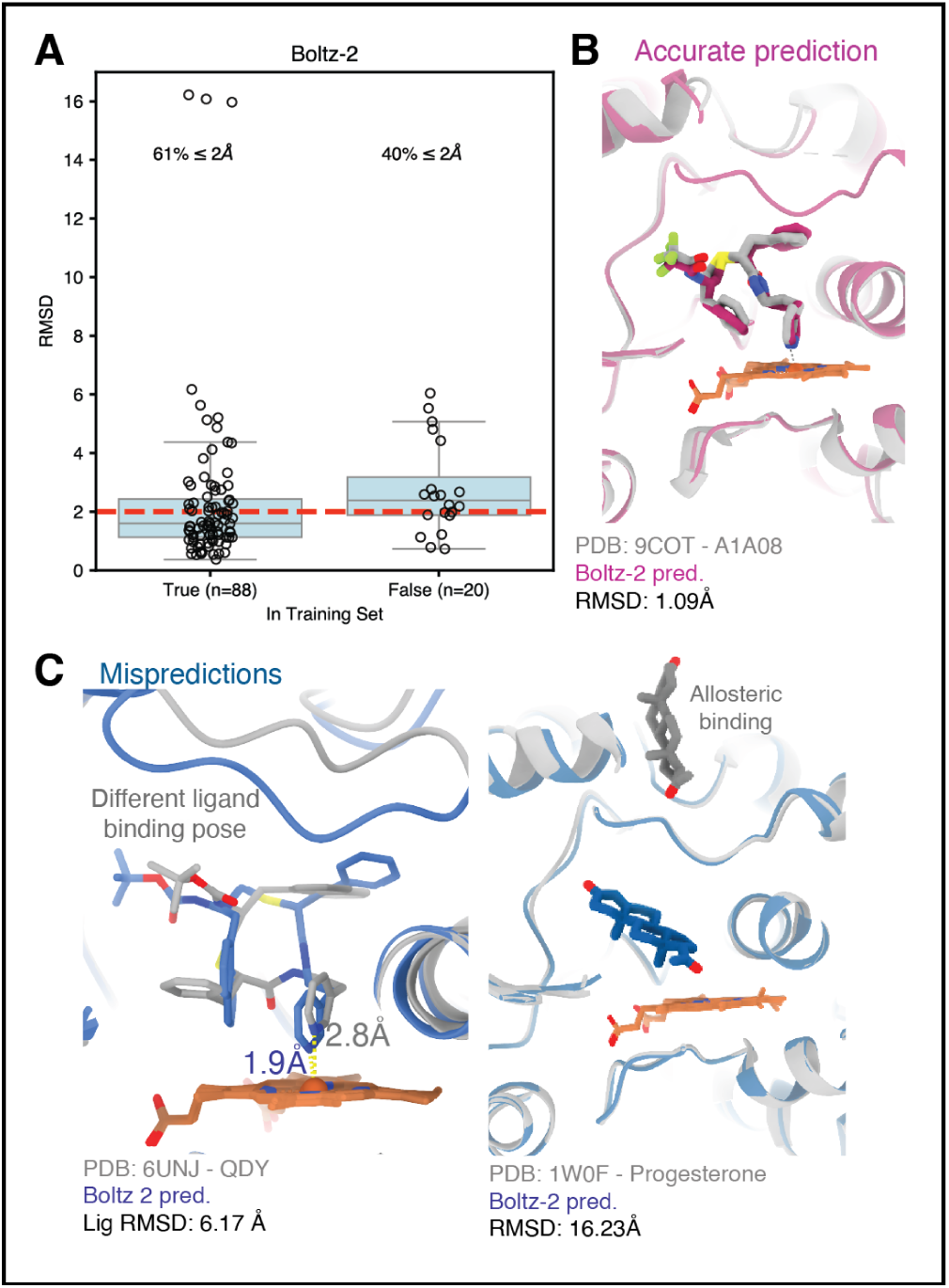
**A.** Overall success rate plot for Boltz-2 predictions. Successful predictions within a ligand RMSD of 2 Å are shown below the red dashed line. The x-axis indicates whether the predictions were before (left) or after (right) the training set cutoff date. **B.** Accurate predictions with a <2 Å ligand RMSD (pink) compared to the crystal structure PDB:9COT (gray). **C.** Mispredictions (blue) with a >2 Å ligand RMSD showing a different binding pose for a ritonavir-like inhibitor (left, PDB:6UNJ in gray) and the wrong binding site of progesterone (right, PDB:1W0F in gray)

### Structural alignments of CYP3A4 reveal regions of coordinated flexibility that can be resolved by cryo-EM

To place the co-folding results and the new cryo-EM structures in the context of the broader CYP3A4 structural landscape, we examined all available CYP3A4 structures in the PDB. One key determinant of binding, the F/G loop, is partially modeled (missing 3 residues or more) or completely unmodeled in 40% of the available structures. The F/G loop is fully modeled in four of the five; only the azamulin complex leaves residues 214–217 unmodeled. Over 95% of structures have unmodeled residues in other regions, most commonly in the loop between helices I and H, likely due to solvent exposure and inherent flexibility (**Fig. 7**A). Outside of this region, loops E/F and G/H are also often partially modeled. This is likely due to the required flexibility of the CYP3A4 active site to expand and allow for large substrates. The five structures reported here follow the same pattern: all show H–I loop disorder, ranging from 7 of 16 residues modeled in the ketoconazole complex to complete continuity in the ritonavir complex, which is the only structure of the five with no chain breaks.

**Figure 7:**
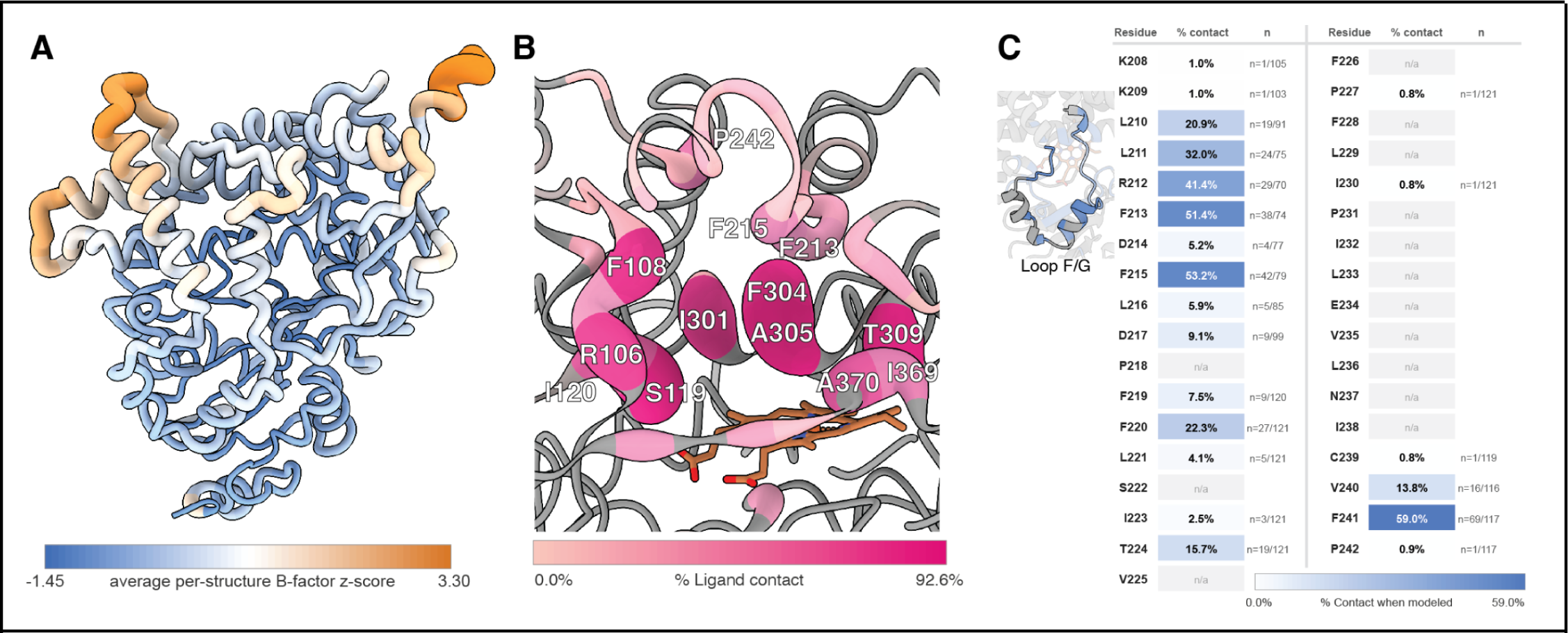
Contact frequency and clustering across 120 CYP3A4 structures. **A.** Worm plot of CYP3A4 colored by the average B-factor z score across all crystal structures in the PDB. Regions with high B-factor values are colored in orange, low B-factor in blue. **B.** Active site residues colored by percent contact occurrence (0-92.6%) across all analyzed structures, displayed on the uninhibited CYP3A4 cryo-EM model. **C.** Isolated statistics of loop F/G on ligand contact when modeled. Inset shows the structure of the helices F’ and G’ along with their connecting loops. Residues that do not form ligand contacts are colored in gray. Contact residues are labeled in a blue gradient. Percentages correspond to the percentage of structures in which a residue contacts the ligand across the structures where the residue is modeled

Across all structures analyzed, contact frequency varied markedly among the 68 active-site residues identified. A core set of residues emerged as near-universal contacts: Ala305 was contacted in 92.6% of structures in which it was modelled, followed by Ser119 (90.9%), Phe304 (86.8%), T309 (80.2%). Ile301 (76.0%) and Phe108 (74.4%)(**Fig. 7**B). As expected, there is a relationship between the ordering of the F/G loop and ligand contact. F/G loop residues Phe215 (53.2%), Phe213 (51.4%), Arg212 (41.4%) show high contact rates with ligands, but are also often excluded from the model due to disorder (**Fig. 7**C). The four ligand complexes reported here recapitulate this hierarchy: all contact Ala305, Ser119, Thr309, and Phe304, and all but ketoconazole contact Phe108. Peripheral engagement scales with ligand size, from 24 contact residues for ritonavir to 17-18 for the smaller ligands. Notably, three of the drugs surveyed in this study showed conformational or compositional heterogeneity which engaged different residues outside of the core set of contacts.

## Discussion

The five cryo-EM structures resolved in this study benefit from the serendipitous C3-symmetric trimeric assembly of a widely used soluble construct. Whether the trimeric arrangement is biologically relevant remains an open question. However, a symmetric trimer does facilitate a historically challenging task of studying CYP enzymes by cryo-EM due to their size. Prior biochemical and biophysical experiments had suggested high orders of oligomeric states were observed in a concentration-dependent manner, with FRET data and crystallographic lattice contacts indicating that substrates such as progesterone could localize to oligomeric interfaces (Davydov et al., 2010). The trimeric state observed here is likely promoted by truncation of the N-terminal transmembrane helix, yet the hydrophobic oligomerization interface maps precisely where the transmembrane helix would be expected, with a putative trimeric helical bundle anchoring the CYPs to the endoplasmic reticulum (**Supplementary Figure 2**). Further investigations with solubilized full-length constructs will be needed to determine whether this interface promotes higher-order oligomerization in the presence of the membrane.

The ability to rapidly screen multiple ligands from relatively small amounts of protein offers a practical advantage when compared to crystallography, as the crystal structures to date have been obtained by co-crystallography and we have had little success with higher-throughput methods such as soaking. The vardenafil result in particular illustrates the value of this approach for ligands without an existing structure, and where computational predictions are insufficient to yield an accurate model. Our work was conducted contemporaneously with work by a contract research organization that also determined cryo-EM structures, yet unreleased in the PDB, of CYP3A4 (NanoimagingServices, 2025), further underscoring the importance of this target for drug development. As detector technology and processing algorithms continue to improve, cryo-EM-based structural characterization of CYP3A4-ligand complexes is likely to become a routine complement to the crystallographic methods that have dominated.

The results here reveal flexibility in the protein consistent with that sampled in previous crystal structures, but also revealing key differences in equivalent structures (e.g. the F/G loop in ritonavir). In addition, we also observe density consistent with multiple conformations (or in the case of ketoconazole, isomers) for multiple ligands, a phenomenon that is likely underreported in the PDB (Flowers et al., 2025; van Zundert et al., 2018). Some of this heterogeneity may be clarified as resolution improves, but the density consistent with multiple binding poses may also be a by-product of the incredible ligand-recognition promiscuity of CYP3A4. As the enzyme most responsible for detoxifying foreign chemicals, it has evolved a binding pocket that can recognize many ligands. Therefore, in addition to flexibility, the poor retrospective performance of co-folding approaches on CYP3A4 ligands and the failure to predict the vardenafil binding mode may also be a consequence of its unusual evolution as a xenobiotic metabolizer.

Overall, the large binding site and the diverse chemotypes recognized by ligands that do not coordinate the heme-iron directly in a Type II mode render structure prediction of CYP3A4 ligand binding poses one of the more challenging problems in early stage drug discovery. These structures emphasize the need for initiatives like OpenADMET (Fraser et al., 2026), where systematically generated, high-resolution structures of avoid-ome anti-targets such as CYP3A4 provide the mechanistic ground truth needed to benchmark and improve co-folding methods. With the new capabilities unlocked by cryo-EM here, such training data can be assembled so that predictive tools can become practically useful in overcoming metabolic issues that arise in drug development.

## Methods

### Δ3-22 CYP3A4 Expression/Purification

A His4- or His6-tagged Δ3-22 CYP3A4 construct was recombinantly expressed in E. coli BL21(DE3) cells co-transformed with the pGro7 chaperone plasmid by heat shock and recovered in SOC medium at 37°C for 40-60 minutes. Transformants were selected on LB agar supplemented with carbenicillin (100 μg/ml) and chloramphenicol (25 μg/ml) overnight at 37°C. A starter culture was grown for 8-14 hours at 37°C in LB supplemented with carbenicillin and chloramphenicol, then used to inoculate 1 L of ZYM-5052 auto-induction medium containing carbenicillin (100 μg/ml), chloramphenicol (25 μg/ml), and 5-aminolevulinic acid hydrochloride (0.5 mM final concentration) to promote heme incorporation. Cultures were grown at 37°C to an OD600 of approximately 0.4, at which point chaperone expression was induced by addition of L-arabinose to a final concentration of 0.5 mg/ml. Temperature was reduced to 25°C and cultures were incubated with shaking for an additional 24 hours. Cells were harvested by centrifugation at 4,000× g for 10 minutes at 4°C and pellets were stored at −80°C.

Cell pellets were resuspended in lysis buffer (50 mM sodium phosphate pH 7.5, 300 mM NaCl, 20% glycerol, 20 mM imidazole, 5 mM beta-mercaptoethanol, 0.5% IGEPAL CA-630) at 100 ml per liter of culture, supplemented with turbonuclease and an EDTA-free protease inhibitor tablet, and stirred at 4°C for 15 minutes. Cells were lysed by sonication on ice at 50% pulse and 50% power for 5 minutes. Cell debris was removed by centrifugation at 35,000×g for 30 minutes at 4°C, and the clarified lysate was filtered and used for purification. Protein was captured by immobilized metal affinity chromatography using a HisTrap HP column pre-equilibrated with load buffer (50 mM sodium phosphate pH 7.5, 300 mM NaCl, 20% glycerol, 20 mM imidazole, 5 mM beta-mercaptoethanol). The column was washed with load buffer and eluted with load buffer supplemented by 300 mM imidazole. The elutions were then further purified by size exclusion chromatography using a HiPrep Superdex 200 column pre-equilibrated with size exclusion buffer (20 mM sodium phosphate pH 7.4, 150 mM NaCl, 10% glycerol, 5 mM beta-mercaptoethanol). Fractions corresponding to the trimeric assembly were pooled, concentrated to 10 mg/ml and flash frozen for storage and cryo-EM sample preparation.

### CYP3A4 ligand binding assay

To probe the binding mechanism of each ligand, 1 μM of purified CYP3A4 was incubated with 50 μM of ritonavir, azamulin, ketoconazole, or vardenafil, to a final volume of 100 μL, and final DMSO concentration of 5%. Triplicate samples along with a control of protein with 5 % DMSO were tested in a 96-well flat bottom black plate (Corning Inc.). The plate was analyzed using a BioTek Epoch 2 Microplate Spectrophotometer by an absorbance measurement in a wavelength scan ranging from 350 nm to 500nm in a 1 nm step. The data was analyzed and plotted using GraphPad Prism 11.0.2.

### Cryo-EM sample preparation and data collection

A key challenge in cryo-EM preparation of CYP3A4 is its dependence on glycerol concentrations of 5% or above for stability, which introduces substantial background noise in vitrified specimens. To mitigate this, protein was purified and concentrated in 10% glycerol, then serially diluted to a final concentration of 3.5 mg/ml in size exclusion buffer without glycerol immediately prior to grid preparation; the final glycerol concentration was 2.5%.

For inhibitor-bound complexes, ligands were dissolved in DMSO to a stock concentration of 100 mM. Protein aliquots were incubated with the ligand at a 10:1 molar ratio, yielding final co-solvent concentrations of 2.5% DMSO and 1.25% ethanol. Complexes were incubated at room temperature for two hours to ensure active site saturation, then diluted to 2.5% glycerol immediately before vitrification.

Quantifoil R1.2/1.3 300-mesh gold grids were glow-discharged for 30 seconds at 15 mA in a Pelco EasiGlow discharge cleaning system prior to sample application. A 3 μl aliquot was applied to each grid and vitrified by plunge-freezing in liquid ethane using a Vitrobot Mark IV (Thermo Fisher Scientific) using Whatman #1 blotting paper with a blot force of 3 and blot time of 3 seconds at 4°C and 100% humidity. To investigate the effect of detergent on preferred orientation, a subset of grids was prepared with 0.08% n-decyl-beta-D-maltopyranoside (DM) added to the sample prior to vitrification. This produced marginal improvement in particle orientation distribution and was not carried forward.

Data was collected on a Talos Arctica 200 keV transmission electron microscope (Thermo Fisher Scientific) equipped with a Falcon 4i direct electron detector. Movies were acquired at a nominal magnification of 190,000x, corresponding to a calibrated pixel size of 0.731 Å/pixel. A total electron exposure of approximately 50 e/Å2 was applied over 40 frames. Data collection was performed using EPU automated acquisition software with a defocus range of −0.8 to −2.0 μm.

### Cryo-EM data processing

All data processing was performed in cryoSPARC. Movie frames were aligned and dose-weighted using patch motion correction and patch CTF estimation. In the unbound dataset, initial particle picking was performed using a template-based picker where the monomeric CYP3A4 (PDB:1TQN) was made into a 3D volume using the molmap tool on ChimeraX 1.11 to a resolution of 5 Å. The volume was then used to generate 10 templates under standard parameters on cryoSPARC, which were then used for template-picker with an expected particle size of 60 Å, yielding approximately 4.3 million particles extracted in a 256-pixel box. Particles were subjected to iterative rounds of 2D classification with standard parameters, 200 classes, and a circular mask of 70 Å to remove junk particles and ice contamination. Inspection of early 2D class averages revealed faint features at the periphery of the protomer, indicating a trimeric complex, which led to the re-extraction of particles at a larger box size of 400px to capture the full particle. Following re-extraction with a 4x bin, 3.9 million particles were subjected to iterative 2D classification as described with 200 2D classes and a 180 Å circular mask, with 937,000 particles selected and carried forward.

Ab-initio reconstruction of 2 volumes with a class similarity of 0 was performed with a starting resolution of 12 Å and a final resolution of 9 Å. This resulted in one trimeric volume and a seemingly monomeric volume. Heterogeneous refinement was then applied to the entire set of 937,000 particles using 3 trimeric volumes and one decoy monomeric volume under standard parameters. Particles assigned to the three trimeric volumes were subjected to an additional round of heterogeneous refinement using the same volumes as the original job. Particles from the three trimeric volumes were then re-extracted under full box size (400 pixels) and subjected to non-uniform refinement imposing C3 symmetry, with an initial resolution of 12 Å. Initial datasets exhibited strong preferred orientation, producing anisotropic maps. Preferred orientation was mitigated by applying 3D classification with 4 volumes and a cutoff resolution of 4 Å, with a solvent mask around the trimeric assembly and a focused mask around a single protomer. Following this procedure, 416,000 particles were carried into a final round of non-uniform refinement with enforced C3 symmetry. Final half-maps were subjected to AR-Decon (Li et al., 2025) to correct for anisotropic resolution, followed by sharpening using EMReady2. Local resolution was estimated using the local resolution tool in cryoSPARC with a mask around the trimeric assembly. Drug-bound CYP3A4 datasets contained a more diverse set of views and therefore AR-Decon was not necessary. In order to better classify the vardenafil occupancy and improve the density, this dataset underwent an additional round of 3D classification for 4 volumes with a focused mask around the active site, without enforced symmetry. The highest resolution map with the best definition of the vardenafil ligand is the output of a Non-Uniform refinement without imposed symmetry. Processing details for the unliganded state and liganded states are shown in (**Supplementary Figure 1**) and (**Supplementary Figure 3**) respectively.

### Model and ligand refinements

Model building was initiated from the CYP3A4 crystal structure PDB:1TQN, which was rigid-body fit into the density and subjected to iterative rounds of real-space refinement in PHENIX v1.21.2-5419 and manual adjustment in Coot 0.9 (Emsley et al., 2010; Liebschner et al., 2019). Our ritonavir, ketoconazole and azamulin structures used reference PDB IDs 3NXU, 2V0M, and 6OOA, respectively. Ligand coordinates and restraints were generated using eLBOW within the PHENIX suite (Moriarty et al., 2009). Model geometry was validated using MolProbity, and refined using Phenix Real Space Refine. Subsequently, the ligands were validated and refined using Schrödinger Maestro (Release 2025-2). The models were prepared through Protein PrepWizard with hydrogen minimization, followed by LigPrep using standard parameters. The protein-ligand model was then subjected to a round of refinement using Phenix/OPLS keeping the protein model as a reference restraint, at a resolution of 3 Å, with a weight of 10, and 2 macrocycles (van Zundert et al., 2021). For models with alternate conformations, the conformations were split into two independent files, refined for ligand pose with fixed protein backbone and side chains, and later merged. One final round of refinement without OPLS restraints was then applied. Figures were prepared using UCSF ChimeraX 1.11.

### Cofolding analysis

Cofolding predictions were generated using the Boltz-2 model, using an automatically generated MSA and predicting five samples for each structure. We excluded any structures from the PDB that had multiple ligands in a single binding site. Using an oracle ranking approach, the sample with the lowest ligand RMSD was selected for reporting, ensuring the results reflect the model’s best possible performance. The ligand RMSD as reported is the ligand heavy atom RMSD after superimposing the binding site and correcting for molecular symmetry (BiSyRMSD as implemented in OpenStructure (Biasini et al., 2013; Robin et al., 2023). This metric provides a complete picture of the reproduction of the ligand’s pose within the binding site. A prediction is considered successful when the ligand RMSD is 2 Å or lower. The training dataset cutoff for Boltz-2 was June 1, 2023.

### Interaction Analysis

The working set comprised 121 single-model PDB-format coordinate files: 116 structures from the RCSB Protein Data Bank and five PDB files representing Azamulin, Ritonavir, Vardenafil, and two poses of Ketoconazole. The chemical identity of each bound ligand was recorded in a companion table (ligand_smiles.csv) that maps each structure identifier to the PDB Chemical Component Dictionary (CCD) code for its ligand and to an isomeric SMILES string for that ligand. For each structure, the first model was read with Biotite, including any CONECT-derived bond information, and the atom array was restricted to chain A. PDB structure 5A1P contained two ligands with CCD codes STR and CIT. This produces a total of 122 protein-ligand pairs.

The receptor was defined as the union of all amino acid residues (identified using Biotite’s amino acid filter) and the heme prosthetic group (residue name HEM). Including the heme in the receptor rather than treating it as a second ligand is essential for CYP analysis: it allows coordination of ligand heteroatoms to the heme iron and van der Waals packing against the porphyrin to be enumerated on the same footing as protein contacts. Intra-receptor connectivity was assigned from residue-name templates, and the resulting structure was converted to an RDKit molecule. Ligand atoms were selected from chain A by CCD residue name and written to an in-memory PDB block, from which an RDKit molecule was created. PDB coordinate records do not contain bond orders, so the correct bond orders, formal charges, and aromaticity were transferred from the reference SMILES for that ligand using RDKit’s AssignBondOrdersFromTemplate.

Protein–ligand interactions were enumerated with a ProLIF fingerprint configured for eleven interaction types: hydrophobic, hydrogen-bond donor, hydrogen-bond acceptor, pi-stacking, anionic, cationic, cation-pi, pi-cation, van der Waals contact, metal donor, and metal acceptor. We used ProLIF’s default geometric criteria (distance and angle cutoffs) throughout. For each structure, the fingerprint was evaluated for every ligand instance against the receptor. The resulting per-residue interaction table was flattened into a list of (receptor residue, interaction type) pairs and annotated with the structure identifier.

We removed hydrophobic contacts at this stage. Because hydrophobic contact is detected for nearly every apolar residue lining the large, promiscuous CYP3A4 active site, keeping it would have filled the matrix with nearly constant features and obscured the directional and polar interactions that distinguish ligands. Van der Waals contacts, detected by the more restrictive ProLIF criterion, were retained. The interactions were collected into a matrix of 122 protein-ligand pairs and 57 residue–interaction features and are provided as CYP3A4_heatmap_data.csv.

All processing was performed in Python 3.11 using Biotite 1.6.0 for structure input and topology perception, RDKit 2026.03.5 for chemical perception and bond-order assignment, ProLIF 2.0.3 for interaction fingerprinting, and pandas 3.0.5 for tabulation.

## Acknowledgements

We thank Irina Sevrioukova (UCI) for the initial construct and expression protocol. A.K.O. is the Rebecca Ridley Kry Fellow of the Damon Runyon Cancer Research Foundation (DRG-2558-25). J.-H.S. is supported by the German Research Foundation (project number 556478029). This work is supported by the Advanced Research Projects Agency for Health (ARPA-H) under AVOID-OME, and Award Number 1AY1AX000035 (to J.S.F.). G.C.L. is supported by the National Institute of Health grant GM143805. The contents are those of the authors. They may not reflect the policies of the Department of Health and Human Services or the U.S. government. The content is solely the responsibility of the authors and does not necessarily represent the official views of the Advanced Research Projects Agency for Health.

## Supplementary Information

**Supplementary Figure 1:**
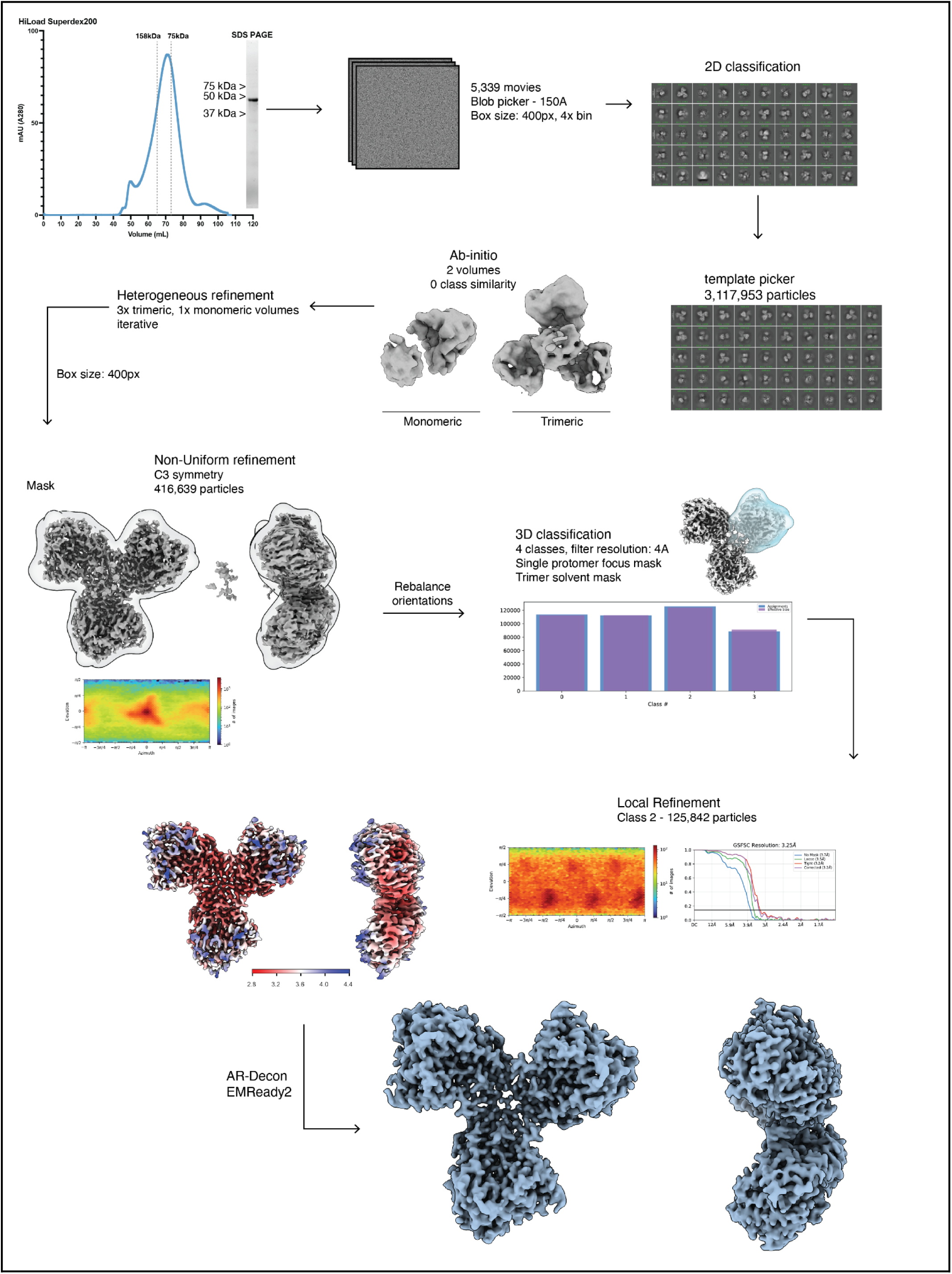
Cryo-EM processing pipeline of CYP3A4. Particles were picked by blob picker followed by 2D classification through cryoSPARC Live. Selected particles were used to generate 2 Ab-initio models which resulted in one volume representing the trimeric state, and a monomeric state used as decoy. After iterative rounds of heterogeneous refinement, 3D classification into 4 volumes sorted the highest resolution into Class 2. The particles were then used for Non-Uniform refinement which resulted in a 3.25 Å map. The final map was sharpened using EMReady2.

**Supplementary Figure 2:**
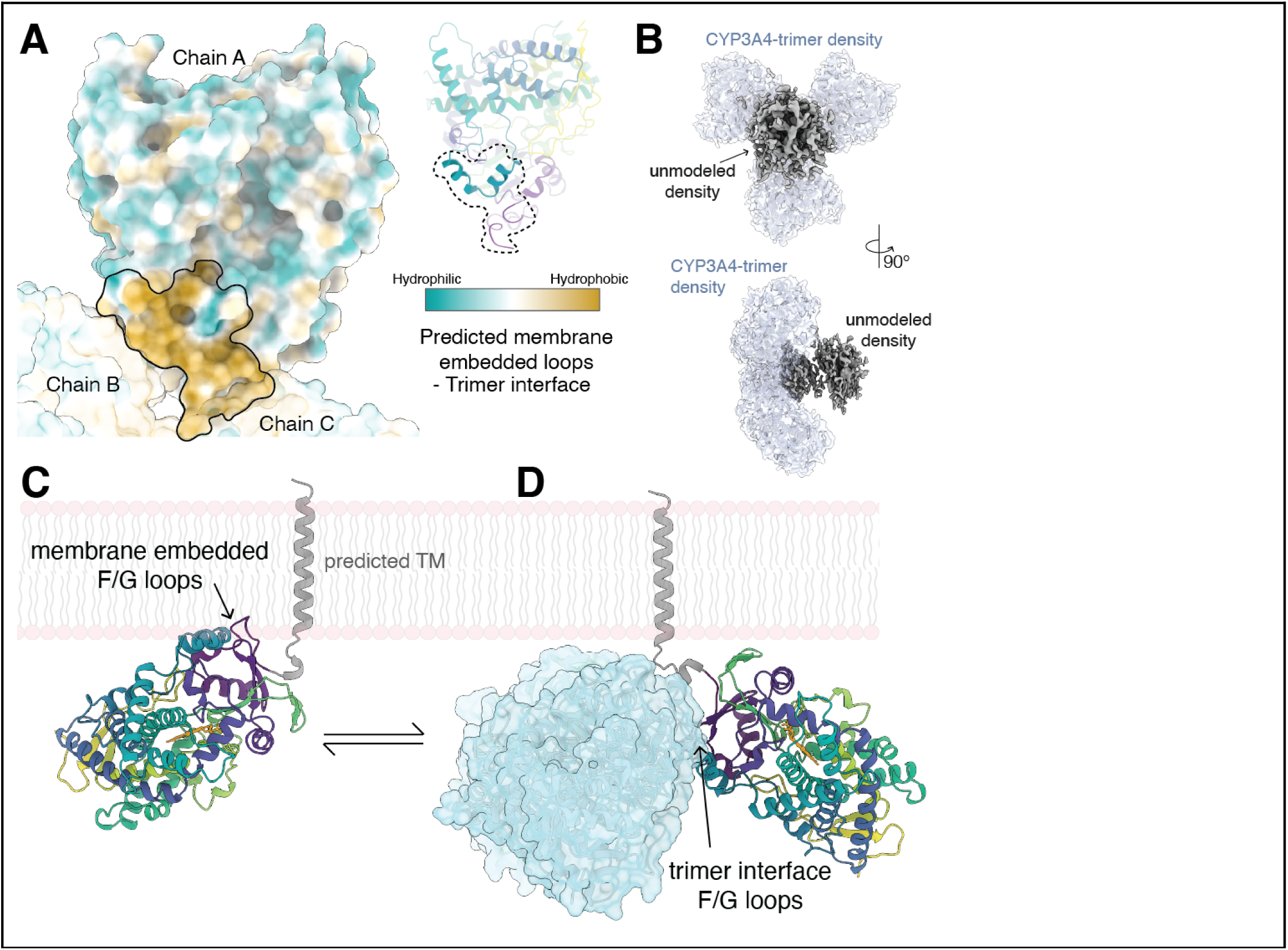
**A**. Surface of a CYP3A4 protomer (left), colored from hydrophilic (teal) to hydrophobic (yellow) by the Kyte-Doolittle scale. The region outlined in black corresponds to the trimer interface, which is predominantly hydrophobic and coincides with the predicted membrane-embedded surface and the trimer interface observed in this study. The corresponding secondary structure is shown at right (viridis), with dashed circles indicating the hydrophobic patch highlighted in the left panel.**B.** Lower contour map of the unliganded CYP3A4, representative of all of the maps in this study, showing unmodeled density (gray). **C**. Monomeric CYP3A4 (shown in the viridis color scheme) in a predicted membrane-embedded configuration, with the F/G loops inserted into the lipid bilayer as predicted by MD (Spanke et al., 2025). The predicted transmembrane helix is shown in gray. **D**. Cryo-EM trimeric assembly, in which the G-helix composes the trimer interface rather than engaging the membrane. We omit the predicted transmembrane helices for the two copies of CYP3A4 represented in cyan surface view. The double arrow indicates the equilibrium between these two states.

**Supplementary Figure 3:**
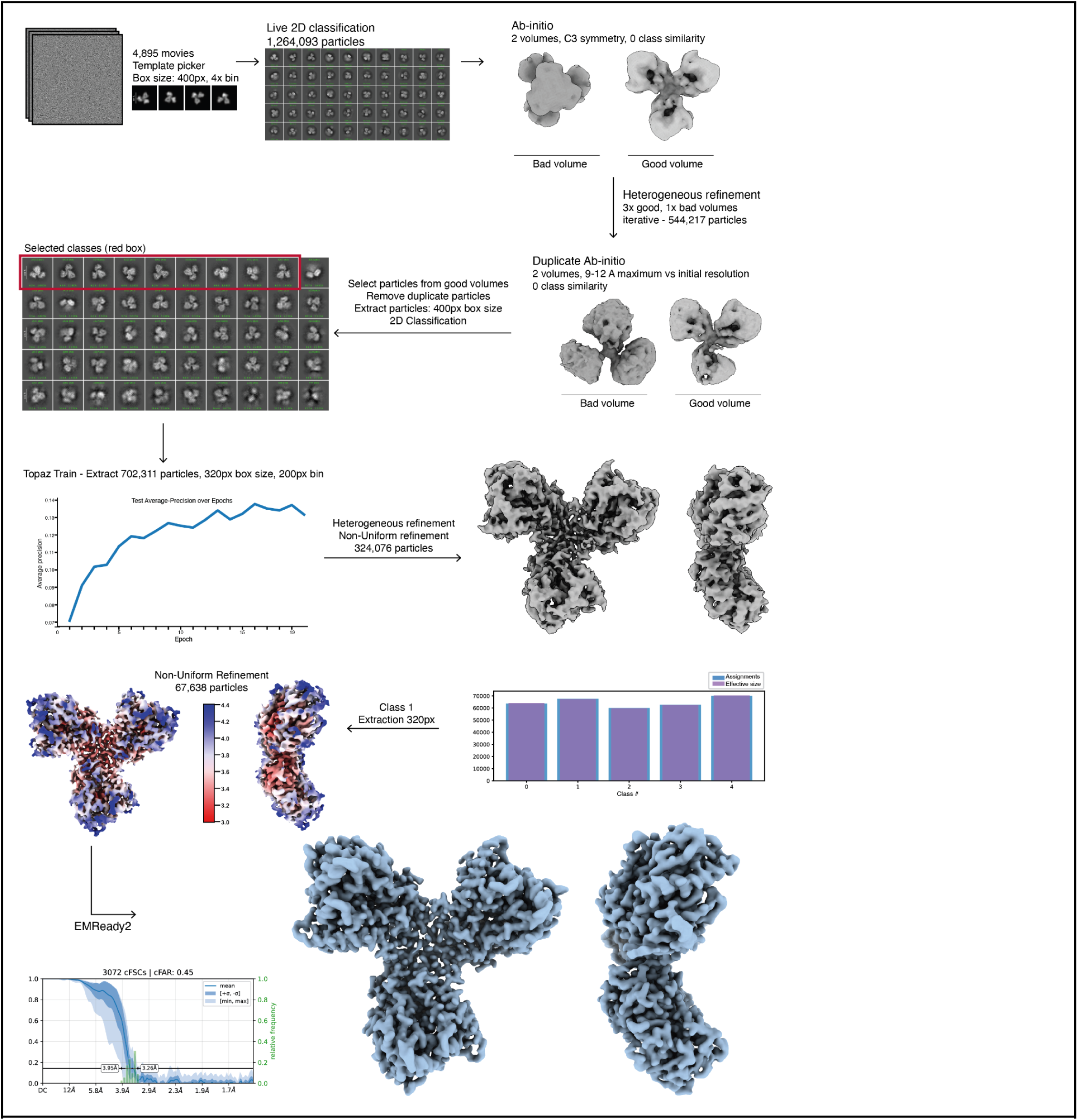
Cryo-EM processing pipeline of ritonavir-bound CYP3A4. Particles were picked using templates from the unbound dataset followed by 2D classification through cryoSPARC Live. Selected particles were used to generate 2 Ab-initio models for heterogeneous refinement. After iterative rounds of refinement, 2D classification was used to select a subset of views for Topaz training. The Topaz picked particles were further refined through heterogeneous refinement. A last round of 3D classification into 4 volumes led to one class with a final reported resolution of 3.55 Å at an FSC of 0.143. The final map was sharpened using EMReady2.

**Supplementary Figure 4:**
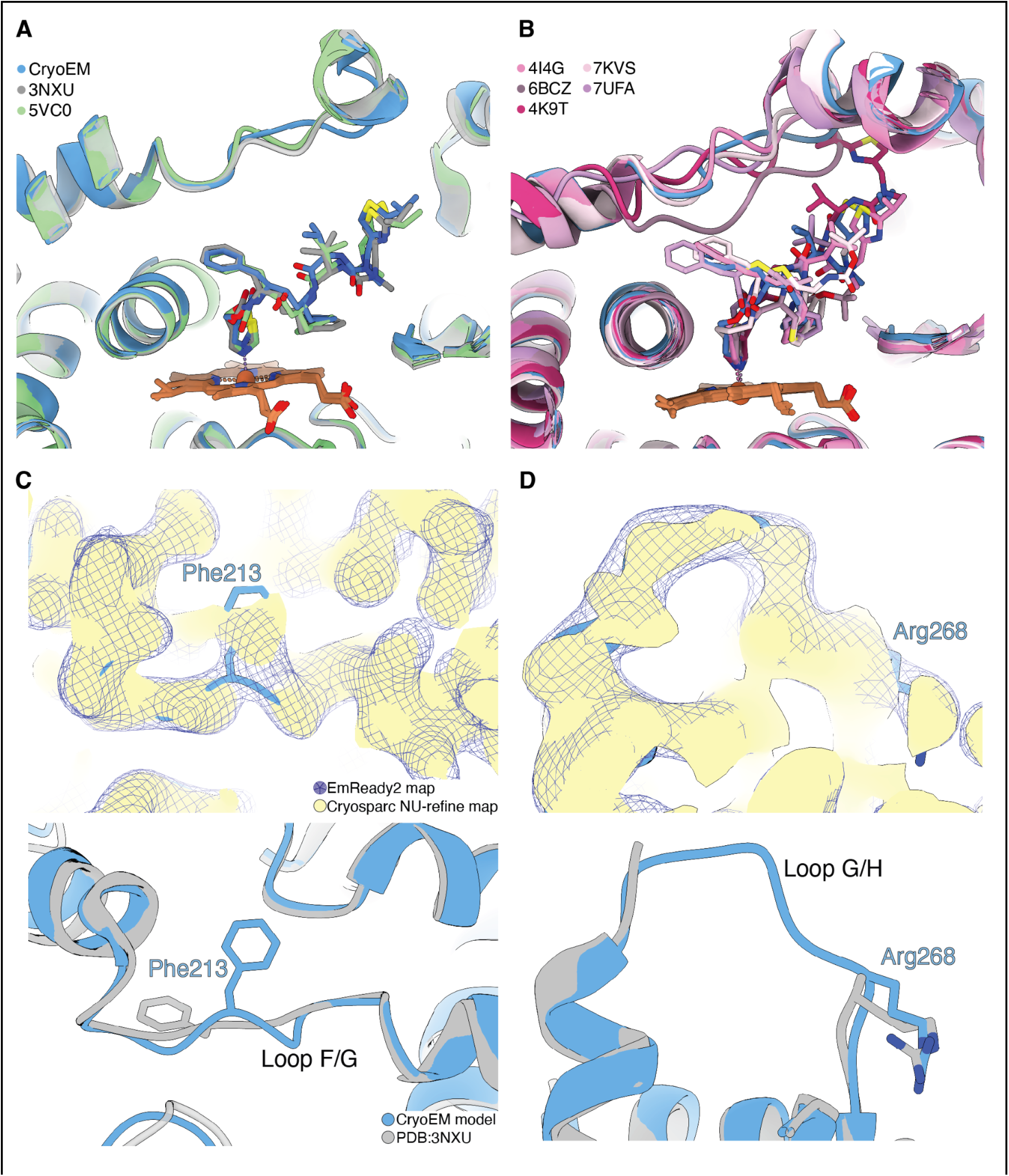
**A.** Overlay of CYP3A4-ritonavir structures PDB: 3NXU (gray), 5VC0 (green) and the cryo-EM model in this study (blue). **B.** Overlay of various ritonavir analogues from diverse publications (shades of pink, PDB: 4I4G, 6BCZ, 4K9T, 7KVS, 7UFA) showing the dynamics of loop F/G in the presence of a highly coordinated ligand like ritonavir and ritonavir-derived molecules. These molecules all share a coordination to the heme iron through a nitrogen-containing heterocyclic ring. **C.** Densities of the F/G loop highlighting residue Phe213. The Non-Uniform refined map from cryoSPARC is shown in yellow, overlaid against the EMReady2 map (mesh) derived from the cryoSPARC map. EMReady2 shows a connected density for the loop along with side chain density for Phe213. This residue is solvent exposed in the cryo-EM model, and coordinates the ligand in the crystal structure PDB:3NXU (bottom panel). **D.** As in **C**, EMReady2 map shows a connected density to residue Arg268, in agreement with the rotameric state in the crystal structure (bottom panel). The loop preceding this residue is unmodeled in PDB:3NXU.

**Supplementary Figure 5:**
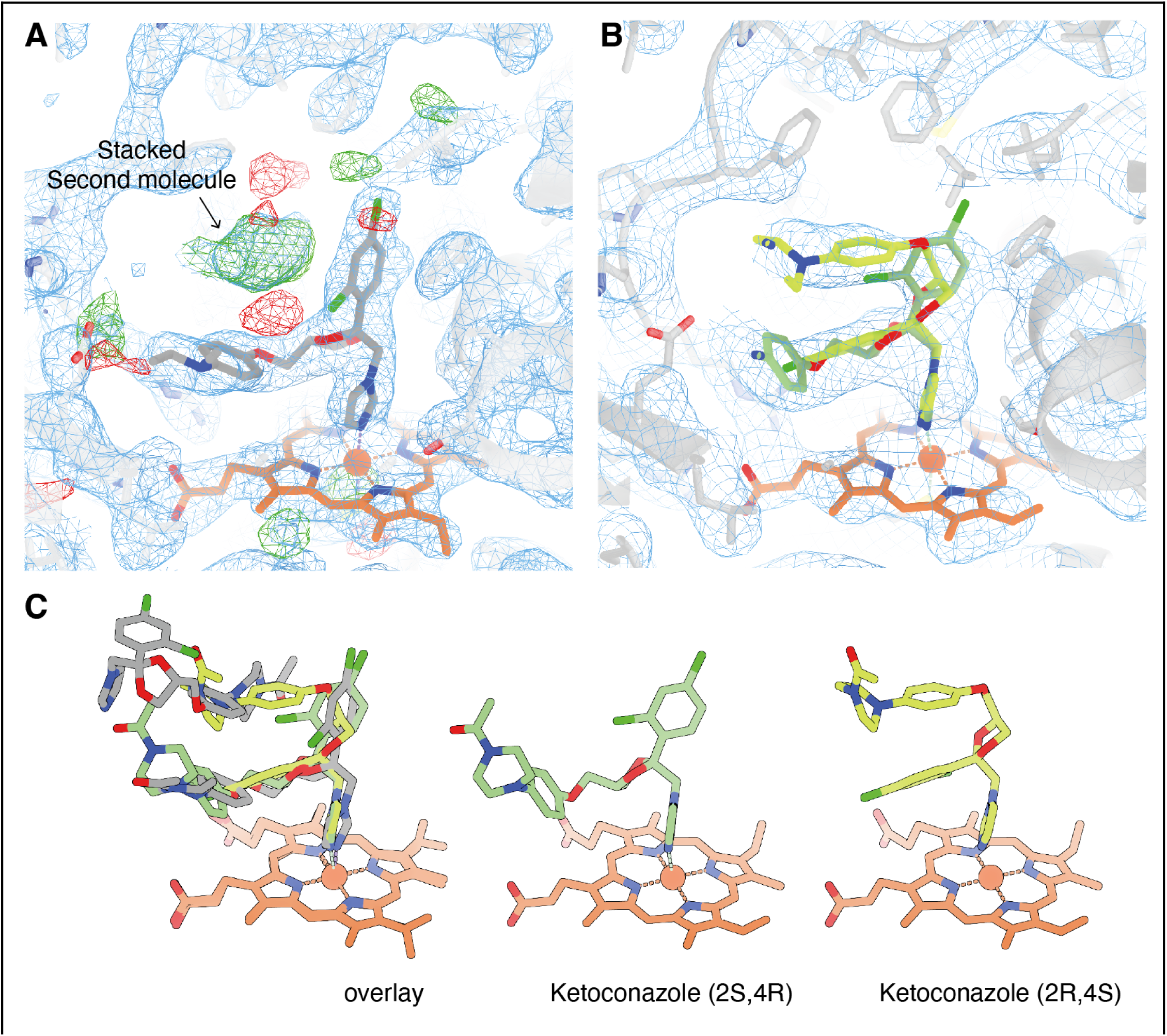
**A.** Re-refined crystallographic map showing difference density (green) for a second stacked ketoconazole molecule, without any connecting density. **B.** cryo-EM density map and model of ketoconazole in two mutually exclusive states with a continuous density between the two molecules. **C.** Atomic view of the ketoconazole molecules coordinating the heme-iron. Left shows an overlay between the crystal (gray) and EM (green) structures. Middle shows the (2S,4R) isomer, right shows the (2R,4S) isomer.

**Supplementary Figure 6:**
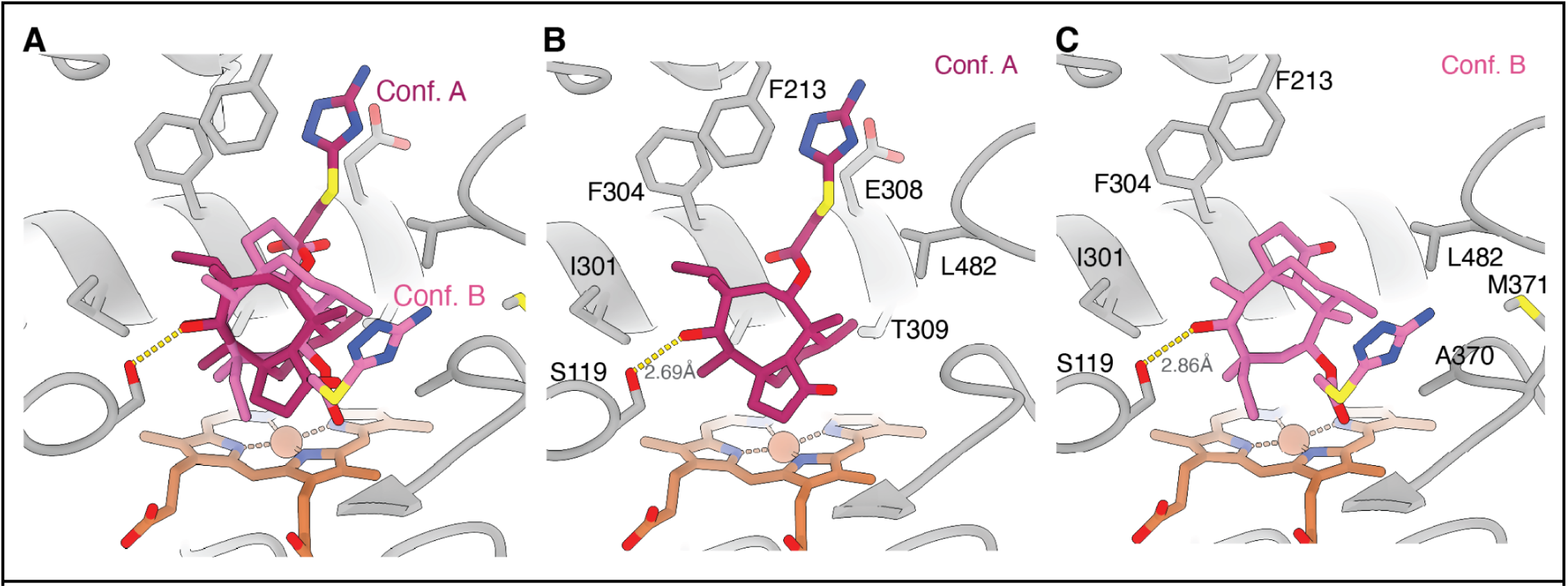
**A.** Conformations of azamulin observed in this study colored in dark (Conf. A) and light (Conf. B) shades of pink. **B-C.** Conformation A and B, respectively, along with the side chain contacts to the CYP3A4 active site.

**Supplementary Figure 7:**
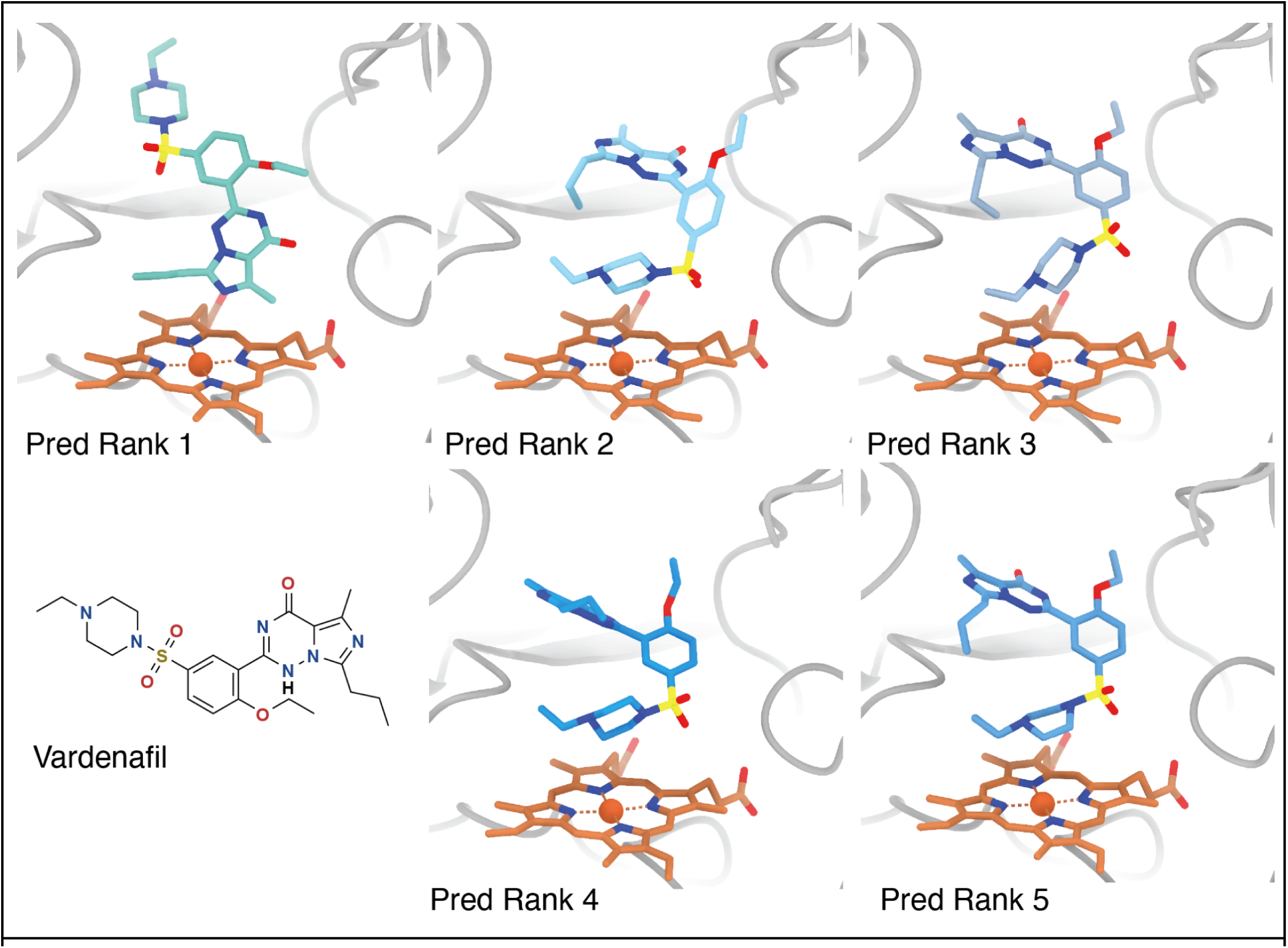
Ranked Boltz-2 predictions of vardenafil-bound CYP3A4. Vardenafil is shown in shades of blue, heme in orange. The chemical structure of vardenafil is shown in the bottom left panel.

**Supplementary Figure 8:**
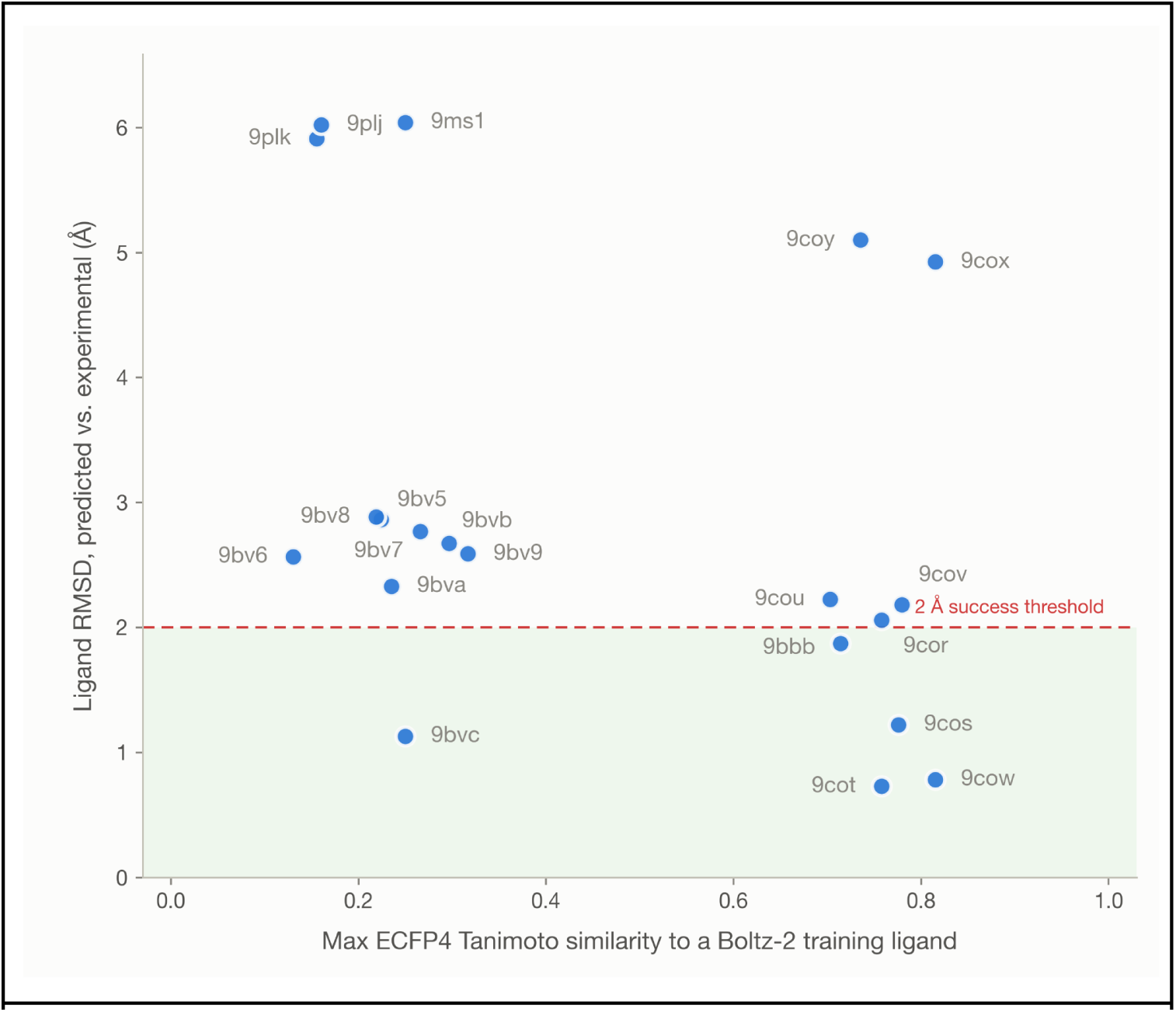
For each held-out structure, maximum ECFP4 Tanimoto similarity to the most similar Boltz-2 training ligand, plotted against ligand RMSD between predicted and experimental pose. The red dashed line marks the 2 Å success threshold.

---

**Table 1.** Atomic modeling, refinement, and validation statistics.

|  | CYP3A4 | CYP3A4 +<br>ketoconazole | CYP3A4 +<br>ritonavir | CYP3A4 +<br>azamulin | CYP3A4 +<br>vardenafil |
| --- | --- | --- | --- | --- | --- |
|  | PDB-38FU<br>EMD-78780 | PDB-38FW<br>EMD-78782 | PDB-38FV<br>EMD-78781 | PDB-38FX<br>EMD-78783 | PDB-38FY<br>EMD-78784 |
| **Model composition** |  |  |  |  |  |
| Protein residues | 1386 | 1344 | 1416 | 1371 | 1422 |
| Ligands | HEM | HEM, KLN, KKK | HEM, RIT | HEM, MWY | HEM,VDN |
| **Model refinement** |  |  |  |  |  |
| Refinement package | Phenix v1.21.2-5419 with OPLS plugin |  |  |  | Phenix<br>v2.2-6143 |
| CC (Volume/mask) | 0.75/0.75 | 0.82/0.83 | 0.73/0.74 | 0.75/0.79 | 0.78/0.81 |
| RMS deviations |  |  |  |  |  |
| Bond lengths | 0.003 (0) | 0.003 (0) | 0.004 (0) | 0.003 (0) | 0.003 (0) |
| Bond angles (°) | 0.700 (0) | 0.632 (0) | 0.704 (0) | 0.702 (0) | 0.621 (0) |
| **Validation** |  |  |  |  |  |
| Map to model FSC 0.5 | 3.5 | 3.6 | 3.7 | 3.6 | 3.5 |
| Ramachandran (%) |  |  |  |  |  |
| Outliers | 0 | 0 | 0 | 0 | 0 |
| Allowed | 1.1 | 1.91 | 1.07 | 1.34 | 1.69 |
| Favored | 98.9 | 98.09 | 98.93 | 98.66 | 98.31 |
| MolProbity score | 1 | 1.11 | 1.02 | 1.25 | 1.18 |
| Poor rotamers (%) | 0 | 0 | 0 | 0.96 | 0 |
| Clash score (all atoms) | 2.24 | 3.14 | 2.42 | 4.82 | 3.95 |
| C-beta deviations | 0 | 0 | 0 | 0 | 0 |
| CaBLAM outliers (%) | 0.44 | 1.65 | 0.86 | 0.46 | 0.21 |
| Q score | 0.416 | 0.351 | 0.423 | 0.393 | 0.478 |

